# Pancreatic tumors activate arginine biosynthesis to adapt to myeloid-driven amino acid stress

**DOI:** 10.1101/2022.06.21.497008

**Authors:** Juan J. Apiz-Saab, Lindsey N. Dzierozynski, Patrick B. Jonker, Zhou Zhu, Riona N. Chen, Moses Oh, Colin Sheehan, Kay F. Macleod, Christopher R. Weber, Alexander Muir

## Abstract

Nutrient stress in the tumor microenvironment requires cancer cells to adopt adaptive metabolic programs to maintain survival and proliferation. Therefore, knowledge of microenvironmental nutrient levels and how cancer cells cope with such nutrition is critical to understand the metabolism underpinning cancer cell biology. Previously, we performed quantitative metabolomics of the interstitial fluid (the local perfusate) of murine pancreatic ductal adenocarcinoma (PDAC) tumors to comprehensively characterize nutrient availability in the microenvironment of these tumors (Sullivan et al., 2019a). Here, we develop <u>T</u>umor <u>I</u>nterstitial <u>F</u>luid <u>M</u>edium (TIFM), a cell culture medium that contains nutrient levels representative of the PDAC microenvironment, enabling study of PDAC metabolism under physiological nutrition. We show that PDAC cells cultured in TIFM, compared to standard laboratory models, adopt a cellular state more similar to PDAC cells in tumors. Further, using the TIFM model we identified arginine biosynthesis as a metabolic adaptation PDAC cells engage to cope with microenvironmental arginine starvation driven by myeloid cells in PDAC tumors. Altogether, these data show that nutrient availability in tumors is an important determinant of cancer cell metabolism and behavior, and cell culture models that incorporate physiological nutrient availability have improved fidelity and enable the discovery of novel cancer metabolic phenotypes.

## Introduction

Altered cellular metabolism is common in cancers (DeBerardinis and Chandel, 2016) and enables many pathological features of tumors (Faubert et al., 2020; Vander Heiden and DeBerardinis, 2017). This has led to substantial interest in identifying the metabolic properties of cancers and the regulation underpinning these metabolic alterations to both delineate the basic biochemistry underlying these diseases and to identify novel therapeutic targets. Recent work has led to the understanding that tumor metabolic phenotypes are driven both by cancer cell-intrinsic factors, such as oncogenic lesions and cellular epigenetic identity (Bi et al., 2018; Nagarajan et al., 2016), and by cell-extrinsic factors in the tumor microenvironment (TME) (Altea-Manzano et al., 2020; Gouirand et al., 2018; Lyssiotis and Kimmelman, 2017; Muir et al., 2018). In contrast to our extensive knowledge of cell-intrinsic regulation of cancer metabolism, how the TME drives altered metabolism in cancers and how such TME-driven metabolic phenotypes contribute to tumorigenesis are poorly understood.

Nutrient availability is a key cell-extrinsic factor that influences cellular metabolism (Elia and Fendt, 2016; Garcia-Bermudez et al., 2020; Muir et al., 2018). Many solid tumors have abnormal vasculature that limits tumor perfusion (Goel et al., 2011; Olive et al., 2009; Provenzano et al., 2012; Wiig and Swartz, 2012), which leads to abnormal nutrient availability in the TME (Gullino et al., 1964; Ho et al., 2015; Vecchio et al., 2021). Thus, perturbed nutrient availability in the TME has been postulated to be a critical driver of cancer metabolic phenotypes (Martínez-Reyes and Chandel, 2021; Reid and Kong, 2013). However, the precise metabolic changes in tumors driven by TME-nutrient cues are largely unknown due to a dearth of information on the nutrient milieu of tumors and a lack of experimentally tractable model systems to study cellular metabolism under such constraints (Ackermann and Tardito, 2019; Cantor, 2019).

To determine how nutrient availability in the TME influences cancer cell metabolism, we recently performed quantitative metabolite profiling of >118 major nutrients and vitamins in the interstitial fluid (the local perfusate of tissues and tumors; IF) in murine models of pancreatic adenocarcinoma (PDAC), providing comprehensive and quantitative knowledge of how TME nutrient availability is altered in PDAC (Sullivan et al., 2019a). Here, we sought to build upon these findings by determining how the abnormal nutrient availability of the PDAC TME drives metabolic phenotypes in PDAC. To do so, we leveraged our knowledge of PDAC IF metabolite concentrations and recent techniques for generating cell culture media with physiological concentrations of nutrients (Cantor et al., 2017; Vande Voorde et al., 2019) to formulate a novel cell culture media termed <u>T</u>umor <u>I</u>nterstitial <u>F</u>luid <u>M</u>edium (TIFM) that recapitulates the nutrient composition of the PDAC TME. This novel cell culture system allows us to study PDAC cells in an experimentally tractable *ex vivo* model while they metabolize substrates at physiological concentrations and identify critical metabolic adaptations of PDAC cells and the TME nutrient cues that drive these adaptations.

To identify such metabolic adaptations, we performed transcriptomic analysis of murine PDAC cells growing in TIFM, standard culture and in orthotopic tumors. Through this analysis, we found that many transcriptional features of PDAC cells growing *in vivo* are better recapitulated in TIFM culture compared to standard culture models. This suggests that altered nutrient availability is a major regulator of the cellular state of cancer cells in the TME and *ex vivo* models incorporating physiological nutrition could improve the fidelity of cell culture models of cancer (Horvath et al., 2016). A major metabolic signature we found in PDAC cells in TIFM and *in vivo* was activation of the amino acid starvation transcriptional signature including increased expression of the *de novo* arginine synthesis pathway. We further find that myeloid driven arginase activity deprives the PDAC TME of arginine and that compensatory activation of the *de novo* arginine synthesis pathway enables PDAC cells in TIFM and in tumors to acquire the arginine needed for amino acid homeostasis despite TME arginine starvation. Collectively, this work identifies TME nutrient availability as a key regulator of the *in vivo* cancer cell phenotype and demonstrates that analysis of cancer cells under physiological nutrient conditions can identify key metabolic features of tumors, such as the activation of *de novo* arginine in PDAC tumors needed to cope with myeloid driven TME arginine starvation.

## Results

### PDAC cells grown in tumor interstitial fluid based culture medium recapitulate the transcriptomic behavior of PDAC cells growing *in vivo*

To study how the nutrient composition of the PDAC TME can influence cancer cell metabolism, we developed a cell culture medium termed <u>T</u>umor <u>I</u>nterstitial <u>F</u>luid <u>M</u>edium (TIFM) based on metabolite concentrations in PDAC IF (Sullivan et al., 2019a) using an approach similar to those described for the generation of media with plasma levels of nutrients (Cantor et al., 2017; Vande Voorde et al., 2019) (Fig. 1A). TIFM is composed of 115 metabolites at the average concentration previously observed in the IF of *LSL-Kras^G12D/+^; Trp53^flox/flox^; Pdx-1-Cre* (KP^-/-^C) (Bardeesy et al., 2006; Sullivan et al., 2019a) murine PDAC tumors. These metabolites were selected on the following bases: (1) commercial availability at high purity, (2) stability in aqueous solution, and (3) presence in pancreatic TIF at a concentration > 0.5µM. To enable rapid identification of bio-active nutrients, TIFM is composed of 9 pools of metabolites that are separately compounded (Cantor et al., 2017; Vande Voorde et al., 2019). To generate the complete medium, the individual metabolite powders are reconstituted in water along with salts at RPMI-1640 (RPMI) concentrations and 10% dialyzed FBS (dFBS) to provide lipids, growth factors and any other macromolecules necessary for cell growth. Sodium bicarbonate is also added at RPMI concentrations to maintain physiological pH (Michl et al., 2019). The complete TIFM formulation is described in Supplementary File 1. Importantly, quantitative metabolite profiling by liquid chromatography-mass spectrometry (LC-MS) of TIFM confirmed that TIFM contained metabolites at expected concentrations, confirming effective compounding of the TIFM metabolite mixture (Fig. 1B). Thus, TIFM recapitulates the nutrient microenvironment of PDAC.

**Figure 1.**
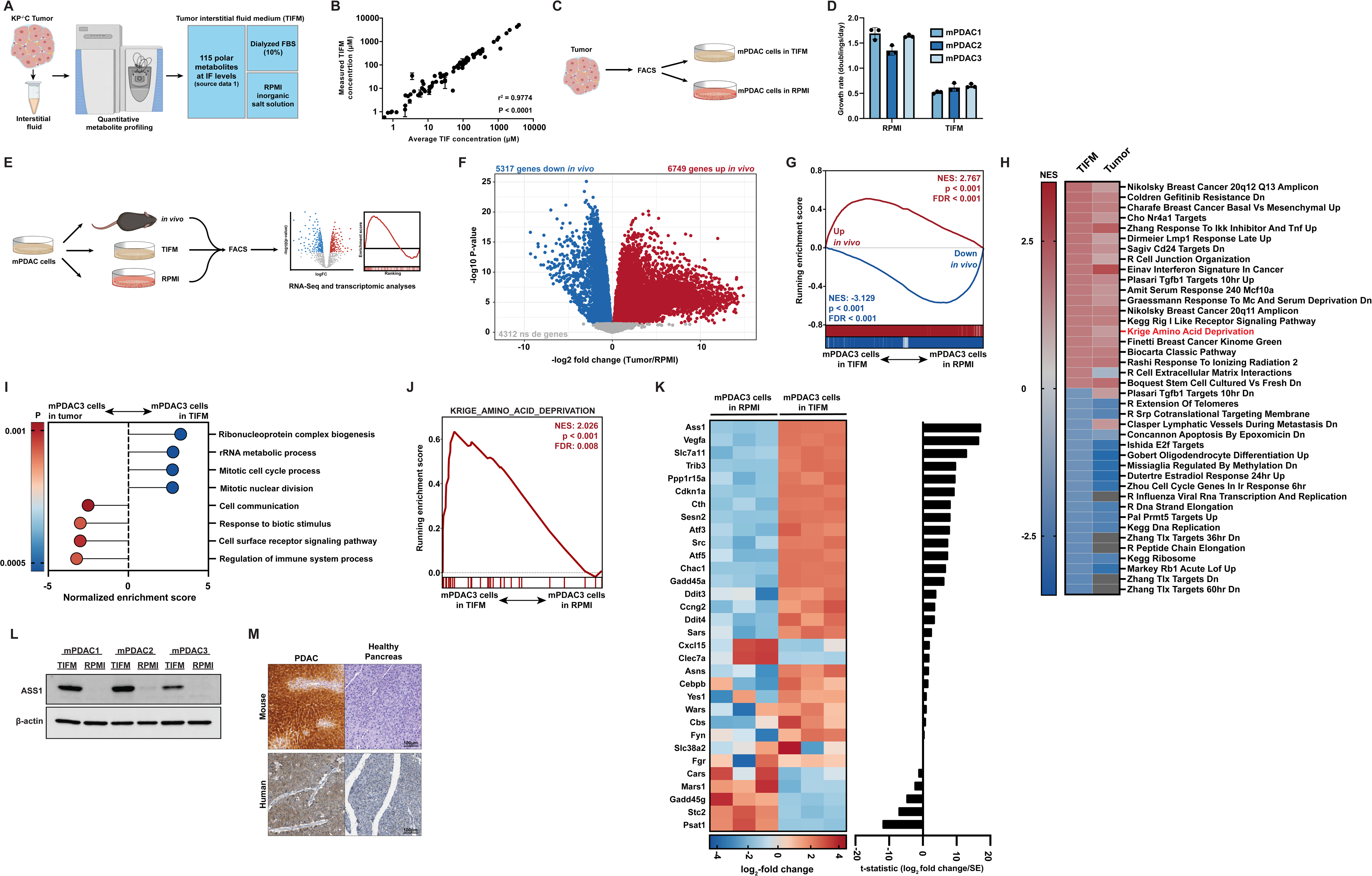
Culturing PDAC cells in physiological nutrients levels better recapitulates the transcriptomic profile of cancer cells in the tumor microenvironment, including changes in expression of arginine metabolism genes. (A) Diagram of the Tumor Interstitial Fluid Medium (TIFM) formulation. (B) Scatter plot of LC-MS measurements of metabolite concentrations in TIFM (n=2) plotted against expected concentrations (average concentration of the given metabolite in KP^-/-^C IF). The values represent the mean of LC-MS measurements and the error bars represent the range. (C) Diagram of the generation of paired PDAC cell lines grown in TIFM and in RPMI isolated from KP^-/-^C tumors, the same tumors used for IF measurements and TIFM formulation. (D) Cell proliferation rate of paired mPDAC cell lines grown in TIFM or RPMI (n=3). The values represent the mean and the error bars represent ± SD. (E) Diagram of workflow for the transcriptomic comparison of mPDAC3-TIFM cells grown in TIFM (n=3), RPMI (n=3) or as orthotopic allograft murine tumors (n=6). mPDAC cells from each condition were isolated by FACS and RNA was isolated for transcriptomic analysis by next generation sequencing. (F) Volcano plot of differentially expressed genes between mPDAC3-TIFM cells in *in vivo* or cultured in RPMI. Blue: downregulated genes with adjusted p<0.05. Red: upregulated genes with adjusted p<0.05. Gray: genes with adjusted p>0.05. Adjusted p-value was calculated through Limma using the Benjamini and Hochberg false discovery rate method (Benjamini and Hochberg, 1995) (G) GSEA of transcriptional differences between mPDAC3-TIFM cells cultured in TIFM or RPMI using custom gene sets from Fig. 1F of those genes up and down *in vivo*. (H) Top 40 enriched (red) or depleted (blue) gene sets from the MsigDB curated gene signature collection mPDAC3-TIFM cells cultured in TIFM compared to culture in RPMI. Also shown are enrichment scores of the same gene sets for mPDAC3-TIFM cells grown *in vivo* compared to culture in RPMI. Grey boxes are gene sets not significantly differentially enriched in mPDAC3-TIFM cells grown *in vivo* compared to cultured RPMI. (I) GSEA analysis using GO gene sets of the transcriptional profile of mPDAC3-TIFM cells *in vivo* compared to mPDAC3-TIFM cells cultured in TIFM. (J) GSEA analysis of the Krige_Amino_Acid_Deprivation signature in mPDAC3-TIFM cells cultured in TIFM or RPMI. (K) Row scaled log_2_ fold change of total gene counts normalized by trimmed mean of M values (TMM) of GSEA plot in Fig. 1J (left) and t-statistic metric for differential expression calculated with limma for these genes (right). (L) Immunoblot analysis of Ass1 in mPDAC cell lines grown in TIFM or RPMI. (M) Immunohistochemical staining for ASS1 in *LSL-KRas^G12D^; LSL-P53^R172H^; Pdx1-Cre* murine PDAC tumors (Hingorani et al., 2005) and healthy murine pancreas (top) as well as in human PDAC tumors and in healthy human pancreas (bottom). Scale bar: 100µm. Human samples are from the Human Protein Atlas (Uhlén et al., 2015).

To determine if TIFM could sustain cancer cells, we isolated murine PDAC (mPDAC) cell lines from three individual KP^-/-^C tumors (the same mouse model used for TIF metabolomics analyses and which formed the basis of TIFM composition (Sullivan et al., 2019a)) by fluorescence activated cell sorting (FACS). We split the isolated cells from each tumor into two populations, which were plated either in TIFM or standard culture conditions (RPMI-1640) to generate paired mPDAC cell lines termed mPDAC-RPMI or mPDAC-TIFM. mPDAC-TIFM cells readily proliferate in TIFM culture, albeit at a slower rate than in RPMI-1640 (Fig. 1D), suggesting that TIFM has the necessary nutrients to sustain PDAC cell proliferation. Interestingly, while mPDAC-TIFM cells continue proliferating when transitioned directly from culture in TIFM to RPMI-1640, transferring mPDAC-RPMI cells directly to TIFM results in complete arrest of cell growth (Fig. 1 – Figure supplement 1), suggesting that long term growth of mPDAC cells in standard cell culture media results in loss of key adaptations to grow under TME nutrient stress. Thus, analysis of PDAC cell metabolism in TIFM could identify novel metabolic adaptations required for growth under TME conditions that would not be apparent from studying PDAC cells under standard culture conditions.

To identify such adaptations, we performed transcriptomic profiling comparing gene expression patterns of the same mPDAC cells (mPDAC3-TIFM) isolated by FACS: (1) after culture in TIFM, (2) after culture in RPMI-1640 and (3) after growing as a syngeneic orthotopic murine tumor to provide an *in vivo* reference (Fig. 1E). This experimental design allowed us to identify transcriptionally-driven metabolic adaptations in TIFM and confirm these were operative *in vivo*. Further, the *in vivo* transcriptomic data allows us to assess how the transcriptional state of PDAC cells in different *ex vivo* models compares to the *bona fide in vivo* cell state. This analysis has recently been suggested to be a critical benchmark for assessing *ex vivo* model fidelity (Raghavan et al., 2021). We first established that compared to standard culture conditions, mPDAC cells in orthotopic tumors substantially alter their transcriptional profile (Fig. 1F and Fig. 1 Source Data 1). In fact, the majority of detected transcripts (12,066/16,378) are differentially expressed in the same mPDAC cells when grown *in vivo* compared to standard culture conditions. Next, we generated sets of genes significantly upregulated and significantly downregulated *in vivo* compared to RPMI. We then performed gene set enrichment analysis (GSEA) (Mootha et al., 2003; Subramanian et al., 2005) on the transcriptional profiles of mPDAC3-TIFM cells growing in TIFM and RPMI using the ‘upregulated *in vivo*’ and ‘downregulated *in vivo*’ gene sets. The gene expression patterns of mPDAC cells growing in TIFM aligns more closely with the expression pattern of mPDAC cells growing *in vivo* (Fig 1G). Smaller gene sets comprised of only the top 500 upregulated and top 500 downregulated genes show the same enrichment patterns with similar enrichment scores confirming there is no enrichment score inflation due to the large gene set size (Fig 1 – Figure supplement 2). We also found a strong correlation between gene expression changes induced by culture in TIFM and growth *in vivo* compared to RPMI (Fig 1 – Figure Supplement 3). Lastly, amongst the top 20 up- and downregulated curated gene signatures from MSigDB (Mootha et al., 2003; Subramanian et al., 2005) in TIFM cultured mPDAC cells compared to RPMI, most where similarly up or down-regulated *in vivo* compared to RPMI (Fig. 1H and Fig. 1 Source Data 2). Together, this transcriptional analysis suggests that mPDAC cells in TIFM better recapitulate the cellular state of mPDAC cells *in vivo*.

We also sought to understand which aspects of the *in vivo* mPDAC cell state were not recapitulated in TIFM. To identify the cellular processes that are differentially regulated between cells growing in TIFM and cells *in vivo*, we performed differential gene expression analysis between mPDAC3-TIFM cells *in vivo* and in TIFM and performed GSEA using Gene ontology (GO) based gene sets. The main cellular processes that differentiate PDAC cells growing *in vivo* from cell growing in TIFM are cell-cell communication, response to biotic stimuli, cell surface receptor activated pathways and regulation of the immune system (Fig. 1I and Fig. 1 Source Data 3). These differences are likely due to the presence of the immune compartment and other neighboring cell populations in PDAC tumors, an aspect of the TME not modeled in TIFM. On the other hand, the main cellular processes positively enriched in PDAC cells in TIFM relative to *in vivo* are ribosome complex biogenesis, rRNA processing and mitotic cell division (Fig. 1I and Fig. 1 Source Data 3), suggesting that, although the slower proliferation of mPDAC cells in TIFM (Fig. 1D) is more reminiscent of cells *in vivo*, cell cycle progression and translation are nevertheless still higher in TIFM than *in vivo*. Altogether, these results show that mPDAC cells grown in TIFM more closely recapitulate the transcriptomic profile of cells growing directly in the TME, suggesting that TIFM is a useful system for the discovery and characterization of cancer cell adaptations to physiological tumor nutrient stress in PDAC.

### Arginine biosynthesis allows PDAC cells to adapt to TME nutrient stress

We next sought to use the transcriptional profiles in mPDAC cells in TIFM and *in vivo,* to identify metabolic adaptations cancer cells exhibit in response to tumor nutrient stress. We focused on adaptation to amino acid deprivation, as this gene signature is one of highly enriched in TIFM (Fig. 1J) and is similarly enriched in mPDAC cells *in vivo* (Fig. 1H). Leading edge analysis (Subramanian et al., 2005) identified *Ass1* as the most differentially expressed gene driving this signature (Fig. 1K). We further confirmed upregulation of ASS1 at the protein level by immunoblotting in TIFM relative to RPMI (Fig. 1L). Immunohistological analysis of murine and human PDAC tumors using data from the Human Protein Atlas (Uhlén et al., 2015) (Fig. 1M) shows similarly robust expression of ASS1, especially when compared to the lack of expression in the healthy exocrine pancreas. These data suggest that mPDAC cell upregulate ASS1 as part of an adaptive response to amino acid deprivation to TME nutrient stress.

Argininosuccinate synthase 1 (ASS1) is the rate-limiting enzyme in the biosynthetic pathway of the non-essential amino acid arginine (Haines et al., 2011). ASS1 catalyzes the synthesis of argininosuccinate from citrulline and aspartate, which can then be converted to arginine and fumarate by argininosuccinate lyase (Asl) (Fig. 2A). Thus, expression of ASS1 enables PDAC cells to synthesize arginine *de novo*. Arginine is one of the most limiting nutrients in the murine PDAC TME at 2-5μM relative to 125μM in plasma, a 20-50 fold decrease (Sullivan et al., 2019a). The TME level of arginine is below the reported Km for arginine transport (Closs et al., 2004). Thus, we hypothesized that mPDAC cells are starved of arginine in the TME and expression of ASS1 allows mPDAC cells to adapt to such starvation by providing an alternative cellular arginine source.

**Figure 2.**
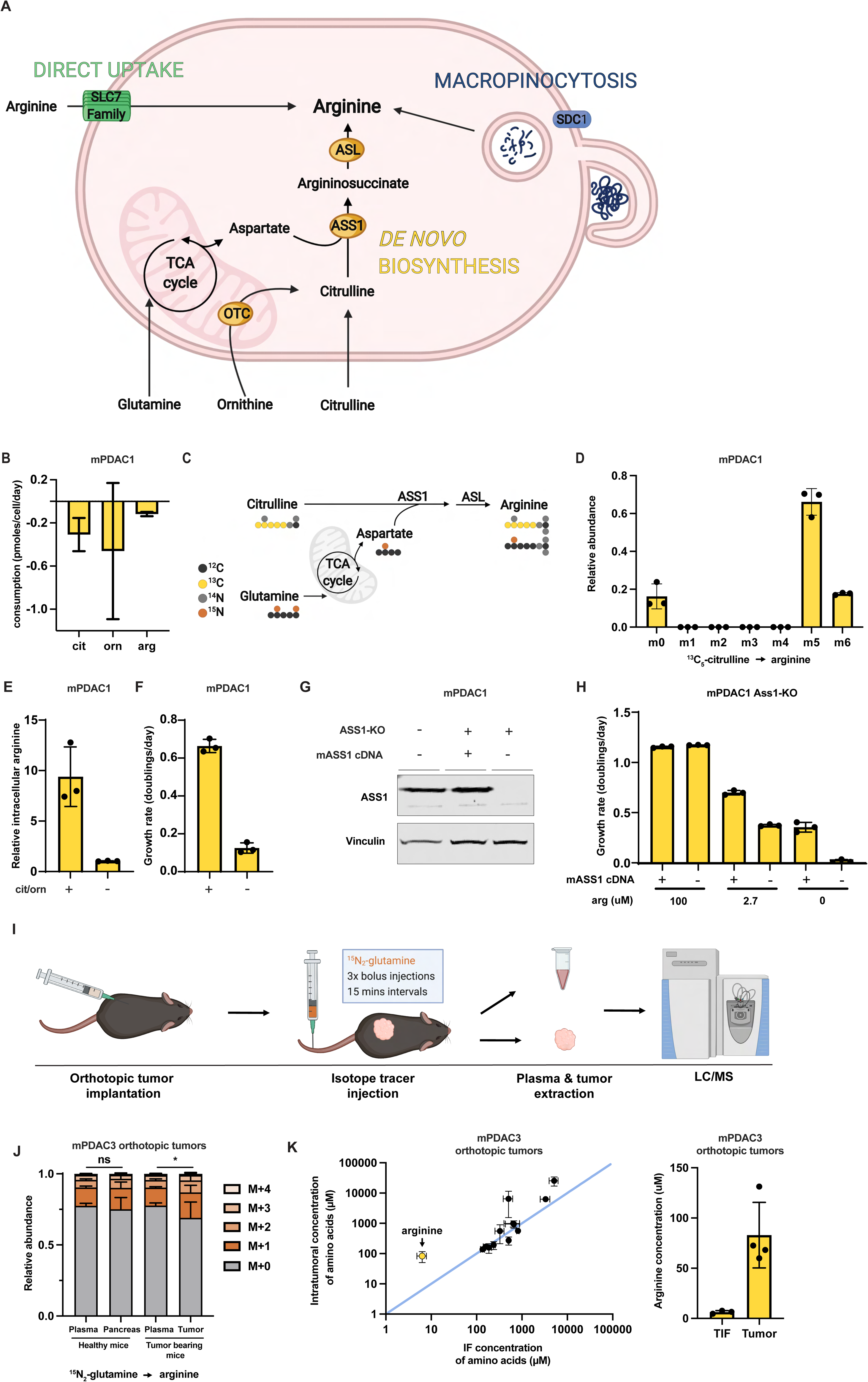
Arginine biosynthesis allows PDAC cells to adapt to low microenvironmental levels of arginine. (A) Cells can acquire arginine by either of three routes: direct uptake of free arginine from the microenvironment, *de novo* synthesis and uptake and breakdown of extracellular protein (macropinocytosis). (B) Cellular consumption/release rate of citrulline, ornithine and arginine by mPDAC1-TIFM cells cultured in TIFM (n=3). (C) Diagram showing cellular metabolic pathways mediating isotopic label incorporation from ^13^C_5_-citrulline and ^15^N_2_-glutamine into arginine. (D) Relative abundance of intracellular arginine isotopomers in mPDAC cells grown in TIFM with ^13^C_5_-citrulline at PDAC IF concentration (67µM) (n=3). (E) Relative intracellular arginine levels of mPDAC cells grown in TIFM with (+) or without (-) TIF concentrations of citrulline (cit) and ornithine (orn). (F) Cell proliferation rate of mPDAC cells in same conditions as (E) (n=3). (G) mPDAC1-TIFM cells were infected with lentiviruses encoding a *Ass1* targeting CRISPR vector. *Ass1* knockout cells were then infected with lentiviruses expressing either CRISPR resistant *Ass1* cDNA or empty vector (E.V.) as indicated. Shown is an immunoblot analysis of these modified cell lines for Ass1 protein expression with vinculin as a loading control. (H) Cell proliferation rate of cells in (G) (n=3). Cells were grown in TIFM with different arginine concentrations as indicated (n=3). (I) Diagram of stable isotope tracing by bolus intravenous injections of ^15^N_2-_glutamine in orthotopic mPDAC3-TIFM tumor bearing mice and non-tumor-bearing controls followed by plasma sampling and tumor extraction for analysis of intratumoral metabolite labeling during the period of kinetic labeling. (J) Relative abundance of ^15^N-labelled arginine isotopomers in tissues or plasma after ^15^N_2_-glutamine tail-vein bolus injections (n=7). (K) (*left*) Concentrations of amino acids in IF (n=3) and tumor samples (n=4) of mPDAC3-RPMI orthotopic tumors measured by LC-MS. (*right*) Bar graph of intratumoral versus IF samples arginine concentrations. For all panels, bar graphs represent mean and error bars represent ± SD. P-value for (G) was calculated using an ordinary one-way anova test and in (K) using a paired, one-tail student’s t test. ns indicates p>0.05, * indicates p<0.05; ** indicates p<0.01; *** indicates p<0.001.

To test if mPDAC cells require *de novo* synthesis to maintain intracellular arginine pools, we first asked if mPDAC cells in TIFM consume the precursors (citrulline or ornithine) used for arginine synthesis. To do so, we used quantitative LC-MS metabolite profiling (Sullivan et al., 2019a) to perform analysis of 108 metabolites that mPDAC1-TIFM and mPDAC1-RPMI cells consume or release in their respective media (Hosios et al., 2016; Jain et al., 2012). We found that mPDAC1-TIFM cells consumed citrulline and ornithine at levels higher than their uptake of arginine (Fig. 2B and Fig. 2 Source Data 1), consistent with active arginine synthesis in TIFM cultured mPDAC cells. We next used stable isotope tracing to measure intracellular arginine derived from *de novo* synthesis. To do so, mPDAC cells were cultured in TIFM with ^13^C_5_-citrulline and steady state incorporation of citrulline carbon into arginine (Fig. 2C) was assessed by gas chromatography-mass spectrometry (GC-MS). The vast majority of intracellular arginine (∼85%) was labeled by the citrulline *de novo* synthesis tracer (Fig. 2D) and this was consistent across multiple mPDAC cell lines in TIFM (Fig. 1 – Figure supplement 1A). Consistent with the citrulline tracing data indicating that *de novo* arginine synthesis contributes the majority of mPDAC arginine in TIFM, inhibiting arginine synthesis by deprivation of the *de novo* synthesis precursors citrulline and ornithine from TIFM results in a 10-fold decrease of intracellular arginine in mPDAC cells (Fig. 2E; Fig. 1 – Figure supplement 1B). This decrease in intracellular arginine is accompanied by a significant decrease in cell proliferation (Fig. 2F; Fig. 1 – Figure supplement 1C). Importantly, individual depletion of either citrulline or ornithine further shows that depletion of citrulline, but not ornithine, is the key substrate mPDAC cells require for arginine synthesis (Fig. 1 – Figure supplement 1D) (Fig. 2B). To confirm the finding that *de novo* arginine synthesis is critical for mPDAC proliferation in TIFM, we used CRISPR-Cas9 to knockout (KO) *Ass1* in mPDAC cells (Fig. 2G). Consistent with decreased mPDAC proliferation upon *de novo* arginine synthesis inhibition by citrulline withdrawal, *Ass1* KO decreases mPDAC proliferation and this affect can be rescued by supplying additional exogenous arginine in TIFM (Fig. 2H). Altogether, these findings suggest that that *de novo* arginine synthesis is important to maintain intracellular arginine levels and mPDAC cell proliferation in TIFM.

We next sought to determine if ASS1-mediated arginine synthesis contributes to arginine homeostasis in murine PDAC tumors. To assess intratumoral PDAC arginine synthesis, we performed ^15^N_2_-glutamine stable isotope tracing by multiple bolus intravenous injections of ^15^N_2_-glutamine into mPDAC orthotopic tumor bearing mice and non-tumor bearing controls (Fig. 2I). ^15^N_2_-glutamine tracing can be used to monitor arginine synthesis and urea cycle activity in PDAC (Zaytouni et al., 2017) by monitoring incorporation of labeled glutamine derived nitrogen into arginine (Fig. 2C). After glutamine injection, healthy pancreas, tumor tissue and plasma samples were collected and ^15^N enrichment in arginine and relevant precursors was measured by LC-MS (Fig. 2 – Source Data 2). Glutamine in plasma is also quickly metabolized by multiple organs and reintroduced into the circulation as intermediate substrates that can contribute to intraorgan and intratumoral labelling (Grima-Reyes et al., 2021). One of the main examples of these interorgan exchange fluxes is the intestinal-renal axis, where glutamine metabolized by the small intestine is released as citrulline and uptaken by the kidneys and further released as arginine for uptake by different tissues (Boelens et al., 2005; Grima-Reyes et al., 2021). Consistent with this, we observed an enrichment of ∼14% ^15^N_1_-arginine and ∼7% ^15^N_2_-arginine in the circulation of healthy and tumor bearing mice (Fig. 2J). We would expect that, for tissues that rely on circulating arginine and not on arginine production, the relative abundance of labelled arginine *in situ* would resemble that of the circulation. In line with this, the relative abundance of arginine in non-ASS1 expressing healthy pancreas resembles the arginine labeling distribution found in circulation (Fig. 2J). In contrast, there is a greater amount of labelled arginine in PDAC tumor tissue compared to plasma, with ∼17% ^15^N_1_-arginine and ∼9% ^15^N_2_-arginine in tumors (Fig. 2J). While these non-steady state isotope labeling experiments cannot allow us to infer the fraction of intratumoral arginine that arises from *de novo* synthesis in PDAC tumors (Buescher et al., 2015), the appearance of additional ^15^N enrichment in intratumoral arginine that cannot be explained by circulating labeled arginine confirms active synthesis of arginine in PDAC tumors, consistent with previous results (Fig. 1) that PDAC tumors highly express ASS1. Lastly, we measured the concentration of amino acids including arginine in the interstitial fluid of orthotopic murine PDAC tumors and compared this to the intratumoral arginine concentration (Fig. 2K and Fig. 2 Source Data 3). We observed that for most amino acids the intratumoral concentration was similar to the IF concentration. However, PDAC tumors had higher concentrations of free arginine than what is present in the TIF. Thus, we conclude that PDAC tumors accumulate higher levels of arginine than available from the local perfusate and that this is at least in part driven by *de novo* synthesis.

### Enhanced uptake of environmental arginine allows PDAC cells to cope with inhibition of *de novo* arginine synthesis

Interestingly, while PDAC cell proliferative capacity is markedly reduced by inhibiting arginine biosynthesis, this doesn’t completely abrogate cell growth (Fig. 2F,H; Fig. 1 – Figure supplement 1C). Given the importance of arginine for cellular homeostasis and growth, we hypothesized that PDAC cells must compensate with other mechanisms to acquire arginine when synthesis is inhibited to maintain viability and proliferation, albeit at a reduced rate (Fig. 2A). In addition to *de novo* synthesis, there are two other known mechanisms for arginine acquisition by PDAC cells: macropinocytosis (Palm, 2019) and cationic amino acid transporter mediated uptake (Closs et al., 2004). Therefore, we sought to determine how these pathways contribute to arginine homeostasis in TIFM cultured mPDAC cells.

To test if macropinocytosis is important for arginine homeostasis in mPDAC cells, we generated mPDAC1-TIFM cells with a doxycycline-inducible shRNA targeting glycoprotein syndecan-1 (SDC1), an important mediator of macropinocytosis in PDAC cells (Yao et al., 2019) (Fig. 1 – Figure supplement 1A). Knockdown of *Sdc1* effectively reduced mPDAC1-TIFM macropinocytosis rate as measured by uptake and catabolism of fluorogenic bovine serum albumin (DQ-BSA), a model macropinocytosis substrate (Fig. 1 – Figure supplement 1B). *Sdc1* knockdown had no effect on intracellular arginine pools nor cell proliferation in TIFM cultured mPDAC cells (Fig. 3A, B). Consistent with this, pharmacological inhibition of lysosomal protein breakdown with hydroxychloroquine (HQ) similarly impairs mPDAC1-TIFM macropinocytosis rate without disrupting cell proliferation (Fig. 1 – Figure supplement 1C, D). Thus, in basal TIFM conditions, macropinocytosis does not appear to be critical for cellular arginine homeostasis. We next tested if macropinocytosis was critical for mPDAC cells upon inhibition of *de novo* arginine synthesis. We measured DQ-BSA uptake and catabolism upon inhibition of arginine synthesis and observed no change in macropinocytosis rate (Fig. 1 – Figure supplement 1E). Furthermore, knockdown of *Sdc1* did not further impair mPDAC cell proliferation upon inhibition of arginine synthesis (Fig. 3C). Thus, we conclude that macropinocytosis does not contribute to mPDAC arginine homeostasis in TIFM, even as an adaptive mechanism upon inhibition of *de novo* arginine synthesis.

**Figure 3.**
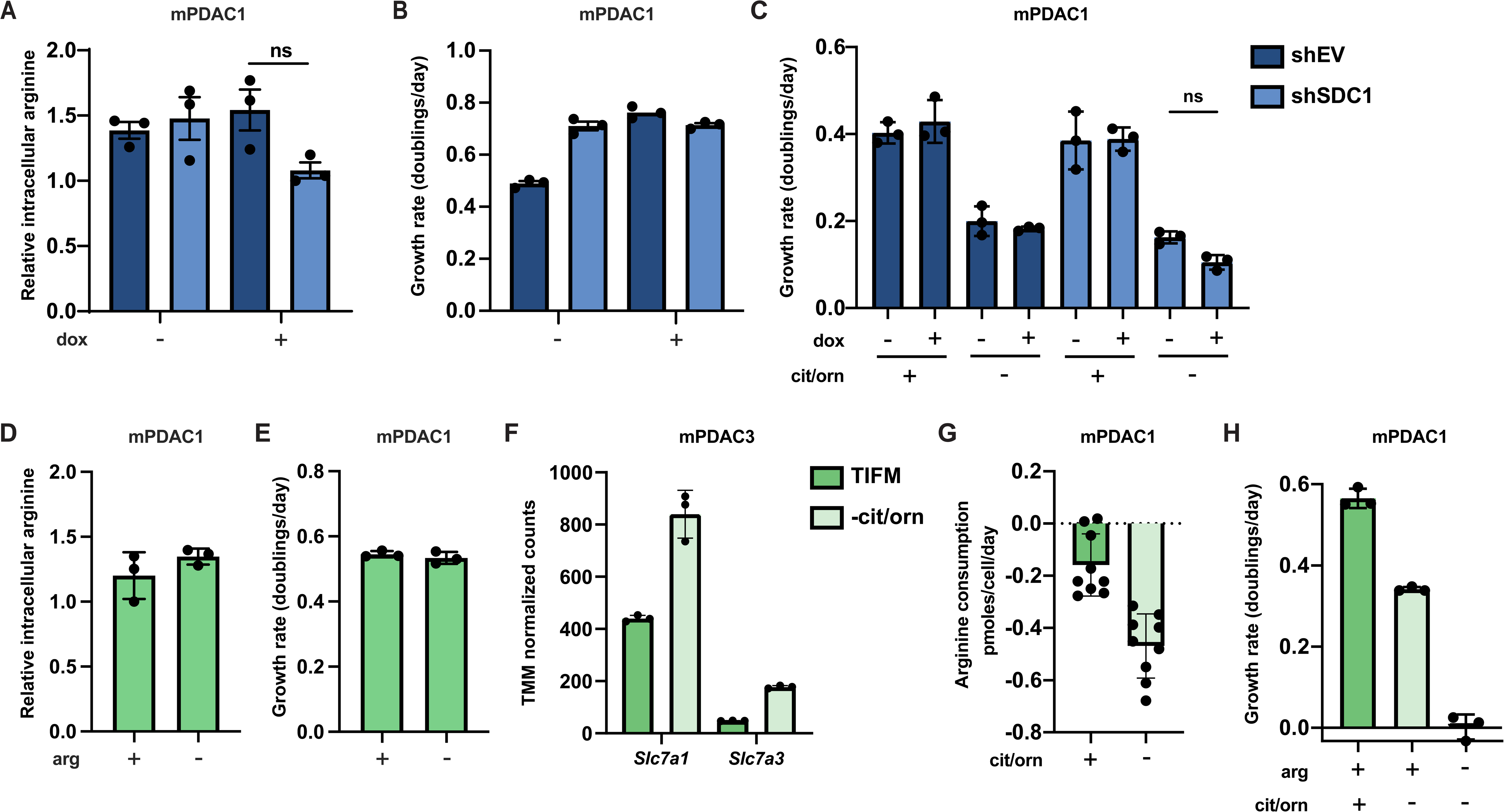
Enhanced uptake of environmental arginine allows PDAC cells to cope with inhibition of *de novo* arginine synthesis. (A) Relative intracellular arginine levels of mPDAC1-TIFM cells infected with lentiviruses encoding a doxycycline inducible *Sdc1* targeting shRNA or an empty vector (EV) control treated with 1 µg/mL doxycycline or vehicle (n=3). (B) Cell proliferation rate of mPDAC1-TIFM cells in same conditions as (A) (n=3). (C) Proliferation rate of mPDAC1-TIFM cells as in (A) cultured in TIFM with or without TIFM concentrations of citrulline (cit) and ornithine (orn) (n=3). (D) Relative intracellular arginine levels of mPDAC1-TIFM cells grown in TIFM with or without TIFM concentrations of arginine (arg) (n=3). (E) Cell proliferation rate of mPDAC1-TIFM cells in same conditions as (D) (n=3). (F) Trimmed mean of M values (TMM)-normalized counts for *Slc7a1* and *Slc7a3,* two cationic amino acid transporters capable of transporting arginine, from transcriptomic analysis (see **Methods**) of mPDAC3-TIFM cells grown in either TIFM or TIFM without citrulline and ornithine (n=3). (G) Per-cell consumption rate of arginine by mPDAC1-TIFM cells cultured in TIFM with or without TIFM concentrations of citrulline and ornithine. Cells were supplemented with 20μM arginine to enable the consumption measurements (n=9). (H) Proliferation rate of mPDAC1-TIFM cells grown with or without TIFM concentrations of citrulline, ornithine, or arginine, as indicated (n=3). For all panels, bar graphs represent the mean and the error bars represent ± SD. P-values for (A) and (C) were calculated using an ordinary one-way anova test and ns indicates p>0.05, * indicates p<0.05; ** indicates p<0.01; *** indicates p<0.001.

We next tested if uptake of the small amount of free arginine in TIFM (∼2.3µM) mediates the ability of mPDAC cells to cope with inhibition of *de novo* arginine synthesis. In normal TIFM culture, removal of arginine does not affect mPDAC intracellular arginine levels nor proliferative capacity (Fig. 3D,E). Thus, as with macropinocytosis, arginine uptake is not critical for mPDAC arginine homeostasis in TIFM conditions. We next tested if depriving mPDAC cells of the available microenvironmental arginine after *de novo* biosynthesis is impaired would affect mPDAC arginine homeostasis and growth. We found that inhibition of arginine synthesis in TIFM cultured mPDAC cells leads to increased transcription of known arginine transporters (Fig. 3F) and leads to an increased rate of arginine uptake by mPDAC cells (Fig. 3G). Furthermore, while we could not detect decreases in mPDAC intracellular arginine levels after eliminating extracellular arginine (Fig. 1 – Figure supplement 1F), eliminating TIFM extracellular arginine completely abrogates cell growth in multiple mPDAC cell lines upon inhibition *of de* novo arginine synthesis (Fig. 3H – Figure supplement 1G). Altogether, these data suggest that mPDAC cells upregulate the uptake of extracellular arginine to cope with inhibition of arginine biosynthesis and that this could be in part mediated by the selective upregulation of cationic amino acid transporters.

### Myeloid arginase causes local arginine depletion in PDAC

Given that PDAC tumors preferentially upregulate arginine *de novo* synthesis to adapt to microenvironmental depletion of arginine, we wanted to understand how arginine becomes depleted in the PDAC TME. Human and murine PDAC tumors contain substantial myeloid compartments (Lee et al., 2021; Zhu et al., 2017) including many arginase-1 expressing cells (Trovato et al., 2019). We confirmed the presence of a robust myeloid and arginase-1 expressing populations in both murine (Fig. 4A) and human PDAC (Fig. 4B) by immunohistochemical analysis. Arginase-1 expressing cells are capable of metabolizing arginine into ornithine and urea (Caldwell et al., 2018). Therefore, we hypothesized that myeloid arginase-1 activity could be responsible for PDAC TME arginine starvation. To test this, we generated orthotopic allograft mPDAC tumors in a mouse model with myeloid specific *Arg1* knockout (*LysM-Cre^+/+-^; Arg1^fl/fl^*) (Clausen et al., 1999; El Kasmi et al., 2008) and control animals (*Arg1^fl/fl^*). We then isolated IF from these tumors at end-stage and measured the levels of amino acids including arginine and ornithine in these samples (Fig. 4C). Compared to control animals, *LysM-Cre^+/+-^; Arg1^fl/fl^* tumors show robust reduction of arginase-1 expression in tumors (Fig. 4D) confirming most arginase-1 in tumors is myeloid in origin. *LysM-Cre^+/+-^; Arg1^fl/fl^* tumors had ∼9-fold increase in IF arginine concentration and a roughly equimolar decrease in ornithine (Fig. 4E and Fig. 4 Source Data 1), consistent with myeloid arginase-1 activity being responsible for TME arginine starvation in these tumors. Pharmacological inhibition of arginase-1 with the small-molecule inhibitor CB-1158 (Steggerda et al., 2017) in mPDAC orthotopic tumors led to a similar increase in IF arginine compared to control treated tumors (Fig. 4F and Fig. 4 Source Data 1). In summary, these results show that arginase activity in the myeloid compartment of PDAC tumors is responsible for arginine depletion in the TME, which results in activation of arginine biosynthesis by PDAC as an adaptation to this metabolic stress in the PDAC TME.

**Figure 4.**
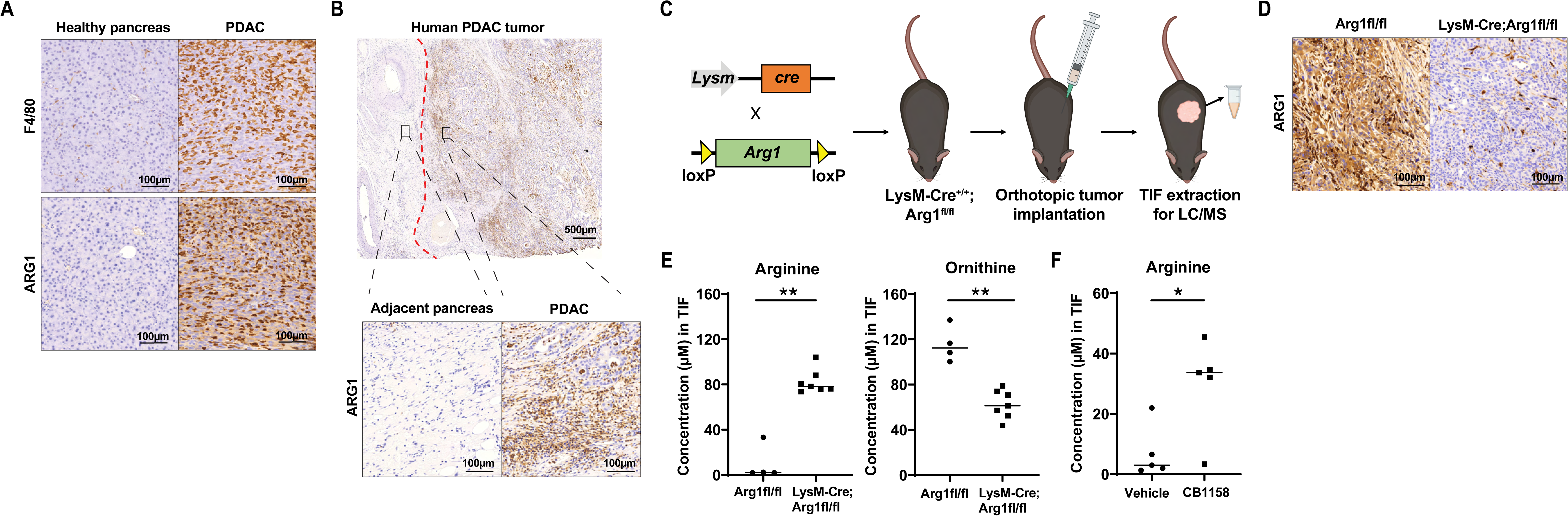
Myeloid arginase causes microenvironmental arginine depletion in PDAC tumors. (A) Immunohistochemical (IHC) staining for F4/80 and ARG1 in an orthotopic mPDAC1-TIFM tumor and in healthy murine pancreas. Scale bars: 100µm. (B) IHC staining for ARG1 in an advanced human PDAC tumor and adjacent untransformed pancreas. Scale bar: 500µm (top) and 100µm (bottom). (C) Schematic for crossing of LysM-Cre and Arg1^fl/fl^, tumor implantation on LysM-Cre^+/+^; Arg1^fl/fl^ progeny and subsequent IF extraction. (D) IHC staining for ARG1 protein expression in orthotopic mPDAC3-TIFM tumors from LysM-Cre^+/+^; Arg1^fl/fl^ and Arg1^fl/fl^ littermate controls. Scale bar: 100µm. (E) Absolute concentration of arginine and ornithine in the IF of orthotopic mPDAC3-TIFM tumors from LysM-Cre^+/+^; Arg1^fl/fl^ (n=7) and Arg1^fl/fl^ littermate controls (n=4). (F) Absolute concentration of arginine and ornithine in the IF of mPDAC3-TIFM orthotopic tumors after treatment with 100mg/kg of arginase inhibitor CB-1158 or vehicle (n=5). P-values for (E) and (F) were calculated using a two-tailed Student’s t test and ns indicates p>0.05, * indicates p<0.05; ** indicates p<0.01; *** indicates p<0.001.

## Discussion

### TME nutrition is a major driver of PDAC cell state

Cell-based models remain critical tools for mechanistic discovery and therapeutic target identification in cancer biology. However, many biological findings and drug targets that arise from these cell-based studies fail to translate to cancer cells *in vivo* or in clinical settings (Horvath et al., 2016). That cancer cells behave so differently in *in vitro* cell-based systems than when in tumors suggests that cancer cell behavior is not completely cell-intrinsically encoded. Rather, cell-extrinsic cues in the TME are capable of dramatically influencing the cancer cell state and impacting many aspects of cancer cell biology including therapy response (Hirata and Sahai, 2017). The importance of TME cues in regulating cancer cell behavior has prompted new efforts to develop cell-based models that incorporate key microenvironmental influences to both improve their disease relevance and fidelity (Horvath et al., 2016) and enable mechanistic studies delineating how the microenvironment influences cancer cell biology.

We directly assessed the impact of the TME on the cellular state of murine PDAC cells by transcriptomic analysis. We found that the TME does indeed induce massive changes in the transcriptional state of PDAC cells compared to PDAC cells in standard culture, consistent with the microenvironment dramatically changing the biology of cancer cells (Fig. 1F). We next specifically asked how TME nutrition influences PDAC cell state and if better recapitulating nutrient access in our cell-based PDAC models would leave PDAC cells in more *in vivo*-like state. Given that metabolism is highly interconnected with epigenetic regulation of gene expression (Chisolm and Weinmann, 2018; Dai et al., 2020; Diehl and Muir, 2020; Reid et al., 2017) and that cellular metabolism is intricately tied to nutrient availability (Elia and Fendt, 2016; Muir et al., 2018), we reasoned that physiological nutrient availability could have dramatic influences on cellular state and be a key microenvironmental factor influencing cancer cell biology. Indeed, we found that growth of PDAC cells in physiological nutrition caused substantial transcriptional reprogramming of PDAC cells that pushed them towards the *in vivo* transcriptional state compared to culture in standard non-physiological conditions (Fig. 1G, H). Thus, consistent with recent studies that have employed cell culture models with physiological nutrient availability (Ackermann and Tardito, 2019; Cantor, 2019; Cantor et al., 2017; Muir et al., 2017; Vande Voorde et al., 2019), we have found that modeling physiological nutrient availability substantially improves cell culture model fidelity. Thus, along with other efforts to improve the fidelity of cell culture models by incorporating microenvironmental factors such as bio-scaffolds enabling three-dimensional growth (Jensen and Teng, 2020; Pampaloni et al., 2007), we anticipate ensuring proper nutrient availability will be critical in the developing more physiologically relevant *ex vivo* cancer models, which will expand our ability to target cancer beyond cell-intrinsic dependencies by allowing the identification and exploitation of microenvironmentally driven therapeutic targets (Metcalf et al., 2021; Sela et al., 2022).

### Arginine is a limiting nutrient in the PDAC microenvironment

Use of the nutrient microenvironment mimicking TIFM model uncovered arginine biosynthesis as a metabolic adaptation that PDAC cells use under microenvironmental nutrient conditions. That PDAC cells would synthesize arginine was initially surprising. Arginine biosynthesis is a metabolically costly process that utilizes intracellular aspartate, a limiting nutrient for tumors (Garcia-Bermudez et al., 2018; Sullivan et al., 2018). Aspartate limitation that arises from arginine synthesis slows nucleotide synthesis and ultimately tumor growth (Rabinovich et al., 2015). Thus, ASS1 acts as a metabolic tumor suppressor and is silenced in many tumor types (Lee et al., 2018). Why would this tumor suppressive metabolic pathway be activated in PDAC?

Multiple studies have also shown that cancer cells can reactivate ASS1 expression and arginine biosynthesis when extracellular arginine becomes limited to support tumor growth. For example, ASS1-silenced tumors treated with arginine deiminase to eliminate extracellular arginine acquire resistance to such therapy by reactivation of ASS1 expression (Rogers and Van Tine, 2019; Rogers et al., 2021). In another example, reactivation of arginine biosynthesis was shown to be necessary to support metastasis of clear cell renal cancers to the arginine limited lung environment, whereas arginine biosynthesis was not necessary and inactive in the arginine-replete primary tumor (Sciacovelli et al., 2021). Lastly, ATF4-CEBPβ mediated upregulation of ASS1 upon amino acid stress has been shown to allow AML cells to adapt to low levels of microenvironmental arginine (Crump et al., 2021). Altogether, these findings suggest that the tumor suppressive role of arginine biosynthesis is context dependent. In the context of microenvironmental arginine deprivation, ASS1 and arginine biosynthesis can switch their role to become tumor supportive. One of the most depleted nutrients in the PDAC TME is the amino acid arginine, which we previously observed was depleted ∼20-50 fold from circulatory concentrations to only 2-5µM (Sullivan et al., 2019a). Thus, given the TME context of PDAC tumors, ASS1 and arginine biosynthesis enables PDAC cells to cope with these TME constraints rather than acting as a tumor suppressor.

Aside from TME arginine restriction in PDAC triggering adaptive responses in the cancer cells, the lack of arginine in the TME can have major impacts on stromal cells that may not have the adaptive capabilities of PDAC cells. For example, anti-tumor lymphocytes require arginine for functionality (Geiger et al., 2016), but are not able to upregulate arginine biosynthesis upon arginine starvation (Crump et al., 2021). Thus, microenvironmental arginine availability is known to limit immune responses in a variety of tumor types (Murray, 2016). This has led to many recent efforts to develop pharmacological tools to increase TME arginine (Canale et al., 2021; Steggerda et al., 2017), which have improved immunotherapeutic outcomes in a variety of murine tumor models (Canale et al., 2021; Miret et al., 2019; Sosnowska et al., 2021; Steggerda et al., 2017). Thus, the severe arginine restriction in the PDAC TME could be a major barrier to immunotherapy in this disease, which is refractory to most immunotherapies (Hilmi et al., 2018). Consistent with this hypothesis, Menjivar and colleagues found that low arginine availability does impair anti-tumor immunity in PDAC and that raising TME arginine levels can improve tumor immune surveillance and response to immunotherapy (Menjivar, co-submitted manuscript). Thus, arginine starvation is a key nutrient limitation that both PDAC and stromal cells face in the TME. Arginine starvation drives PDAC metabolic adaptation but other cell types without adaptive capacity, such as lymphocytes, face arginine starvation driven dysfunction in the TME. Future studies delineating how different cellular populations are affected by TME arginine starvation will prove critical to better understanding how tumor physiology impacts cancer and stromal cell biology.

### Stromal control of microenvironmental nutrient conditions

We previously had found that the TME is arginine depleted (Sullivan et al., 2019a). However, what drove arginine depletion in the TME was unknown. Here, we find that the large arginase-1 expressing myeloid compartment in PDAC tumors is largely responsible for TME arginine depletion (Fig. 4). Consistent with these findings, Menjivar and colleagues also found PDAC associated myeloid cells are critical for mediating TME arginine depletion (Menjivar, co-submitted manuscript). Thus, the most striking nutrient perturbation in the TME is not driven by abnormal cancer cell metabolism but is instead driven by stromal metabolic activity. This finding is in line with recent studies documenting the critical role that stromal cells have in influencing nutrient availability in the TME. For example, in addition to the role we have found for myeloid cells in limiting TME arginine, myeloid cells were found to be the major glucose consuming cell type in a variety of tumor types (Reinfeld et al., 2021). Thus, stromal myeloid cells are likely key regulators of glucose availability in the TME. Fibroblasts have been show to also regulate levels of key metabolites in the TME (Sherman et al., 2017), such as amino acids (Francescone et al., 2020; Sousa et al., 2016) and lipids (Auciello et al., 2019). Recently, tumor innervating neurons were also shown to regulate availability of amino acids in the TME (Banh et al., 2020). Thus, future studies delineating the complex metabolic interactions amongst tumor and stromal cells (Li and Simon, 2020) will be critical to understanding how nutrient availability is regulated in the tumor ecosystem and the resulting nutrient milieu impacts cancer and stromal cell metabolism and biology.

## Figure Legends

**Figure 1 – Figure supplement 1.**
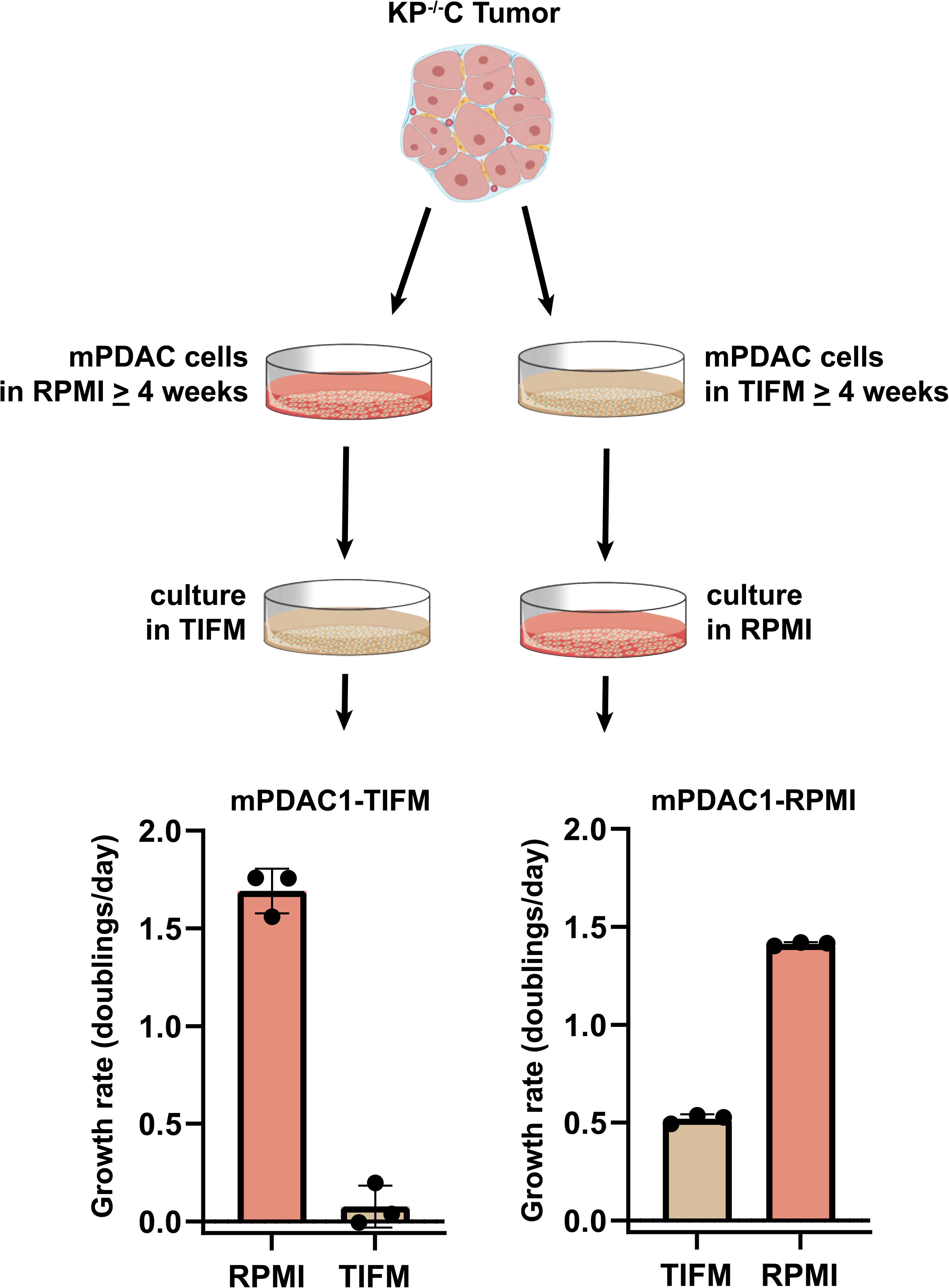
mPDAC cells cannot proliferate in TIFM after long term culture in RPMI. mPDAC cells were isolated from a KP^-/-^C tumor and cultured in either TIFM or RPMI directly. After long term culture (>1 month) in either media condition, cells were passaged into the other media (i.e. cells grown in TIFM were subsequently cultured in RPMI and cells grown to RPMI were subsequently grown in TIFM). The proliferation rate after each media swap was measured (n=3). The values represent the mean and the error bars represent ± SD.

**Figure 1 – Figure supplement 2.**
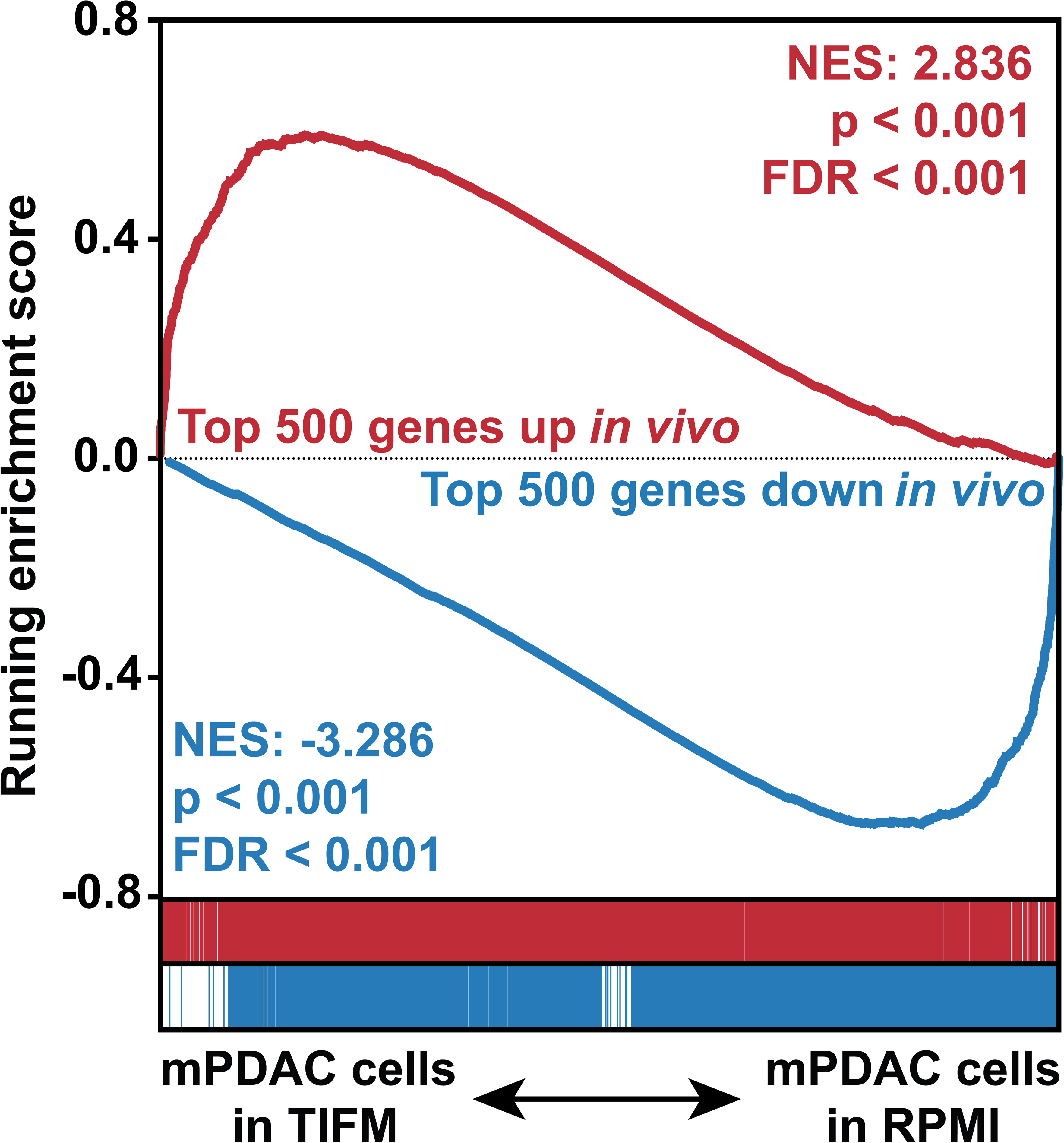
GSEA using gene signatures of the top 500 upregulated and top 500 downregulated genes *in vivo* vs. RPMI shows TIFM grown cells better recapitulate the transcriptomic behavior of PDAC cells *in vivo*. Custom gene sets of the top 500 genes up- and downregulated (by adjusted p-value) in mPDAC3-TIFM cells growing *in vivo* compared to standard culture were generated from the data in Fig. 1E,F. GSEA analysis was performed on transcriptional data of mPDAC3-TIFM cells cultured in TIFM and RPMI using these gene sets.

**Figure 1 – Figure supplement 3.**
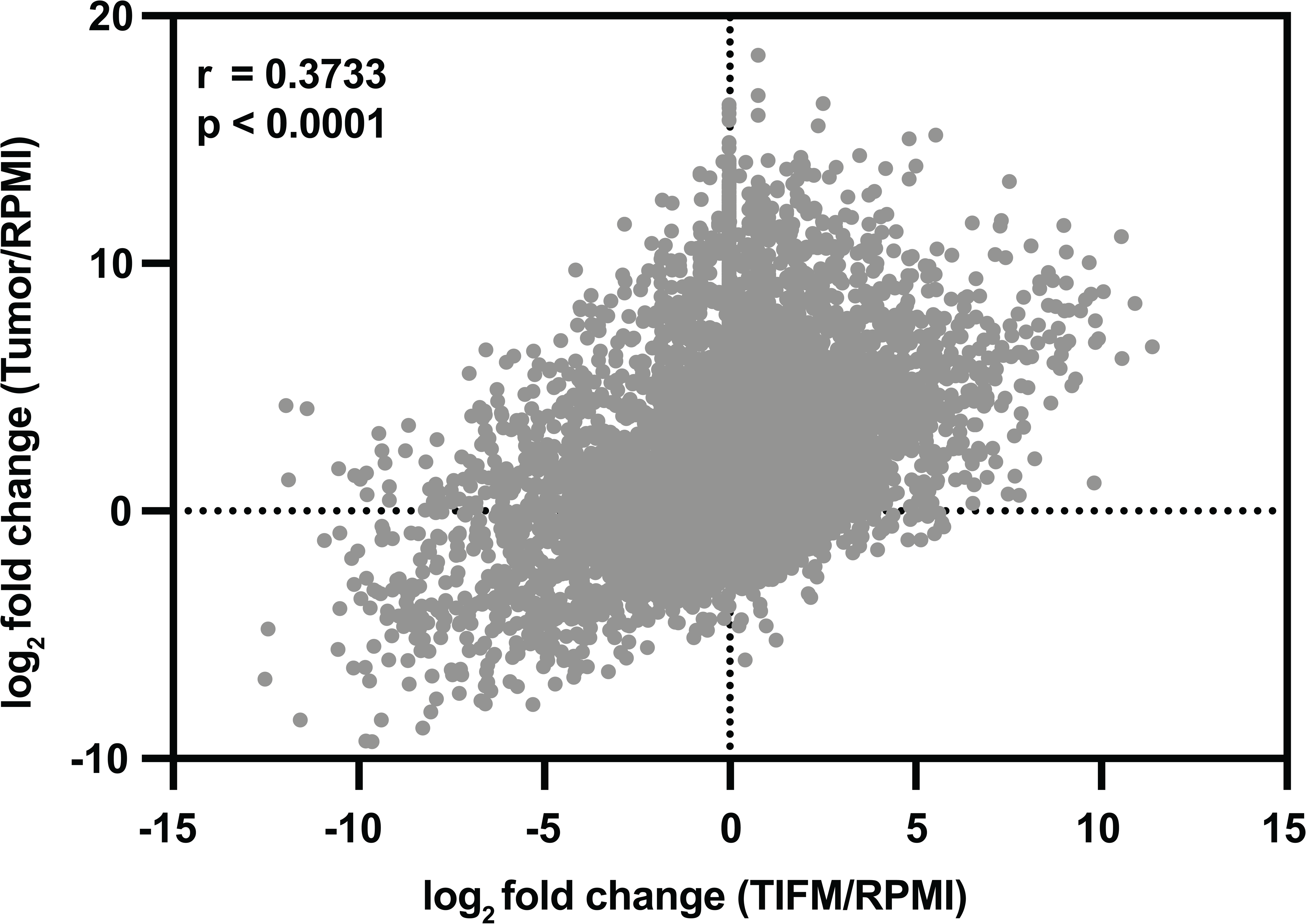
Correlation between gene expression changes in mPDAC3-TIFM cells cultured in TIFM and *in vivo*. Mean log_2_ fold changes in gene expression between mPDAC3-TIFM cells cultured in TIFM (n=3) versus RPMI standard culture reference (n=3) and compared to mPDAC3-TIFM cells isolated from an orthotopic tumor (n=6) compared to RPMI. Pearson correlation r = 0.3733, p<0.0001.

**Figure 2—Figure supplement 1.**
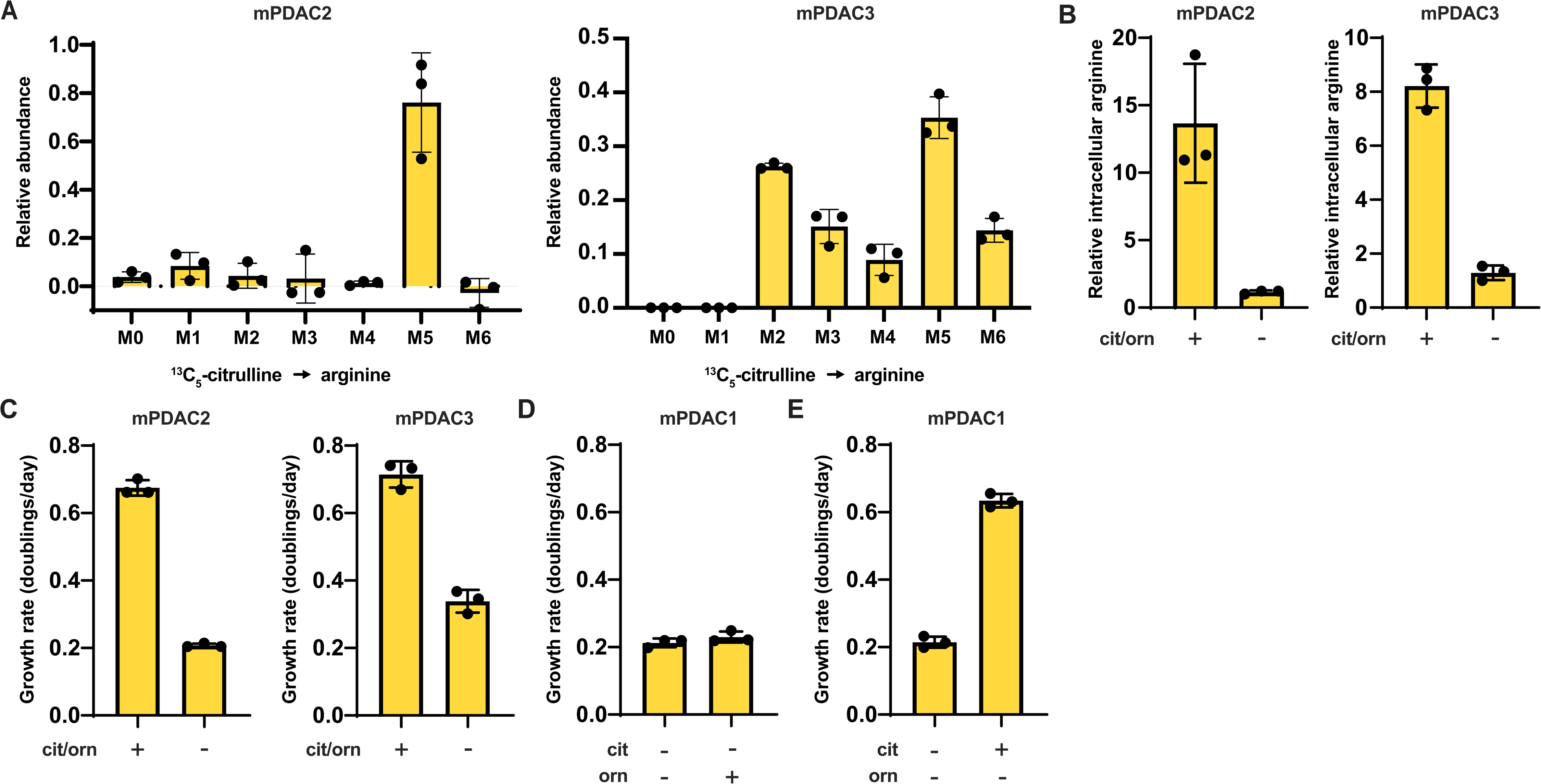
Arginine biosynthesis allows for adaptation to low physiological levels of microenvironmental arginine in multiple mPDAC cell lines. (A) Relative abundance of intracellular ^13^C-labelled arginine isotopomers in mPDAC2-TIFM and mPDAC3-TIFM cells grown in TIFM with ^13^C_5_-citrulline (n=3). (B) Relative intracellular arginine levels of mPDAC2-TIFM and mPDAC3-TIFM cells grown in TIFM with (+) or without (-) PDAC IF concentrations of citrulline (cit) and ornithine (orn) (n=3). (C) Cell proliferation rate of mPDAC2-TIFM and mPDAC3-TIFM cells in same conditions as (B) (n=3). (D) Cell proliferation rate of mPDAC1-TIFM cells with or without TIFM concentrations of ornithine (orn) (n=3). (E) Cell proliferation rate of mPDAC1-TIFM cells with or without TIFM concentrations of citrulline (cit) (n=3). For all panels, bars represent the mean and error bars represent the mean and the error bars represent ± SD.

**Figure 3—Figure supplement 1.**
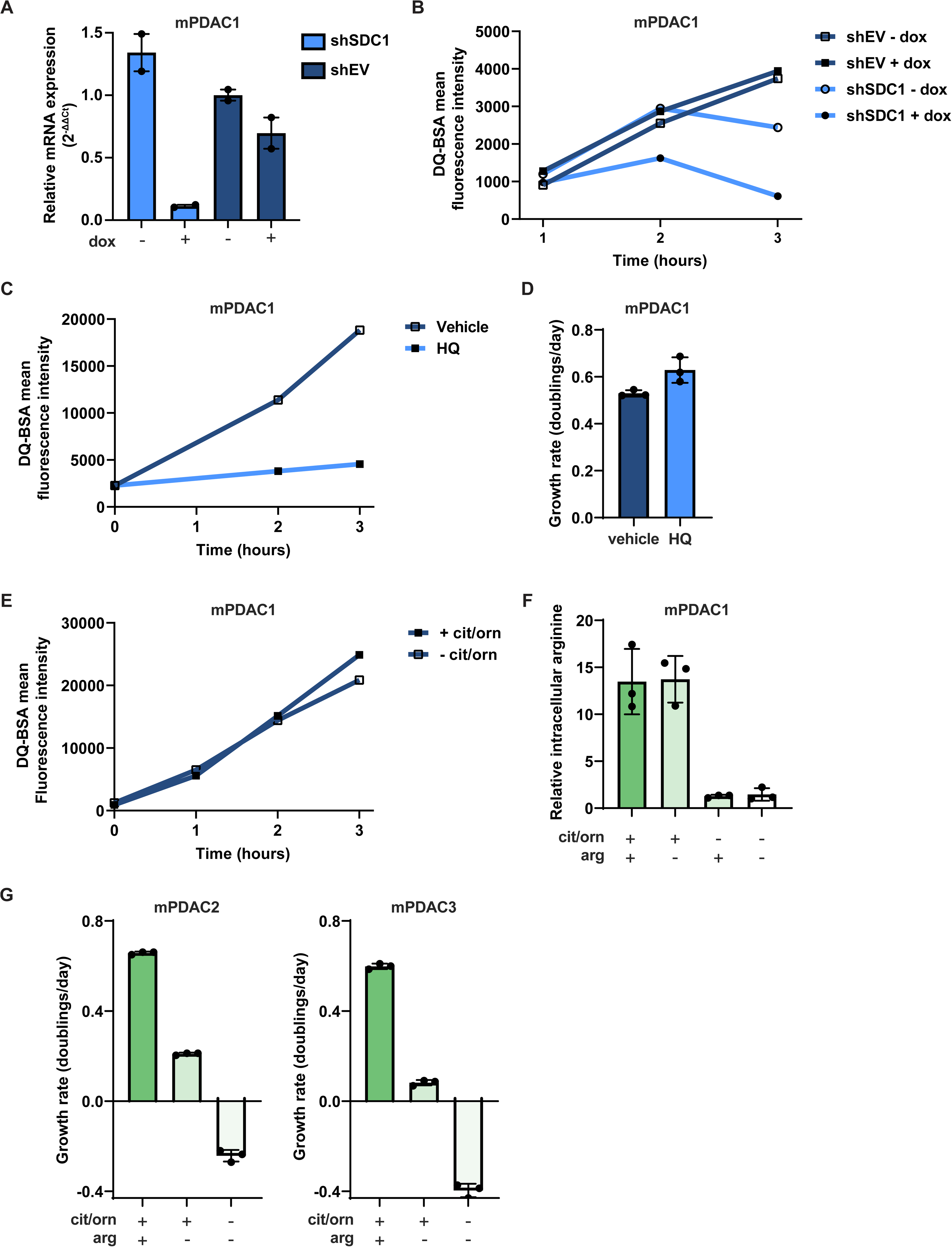
mPDAC cells do not upregulate macropinocytosis after inhibition of arginine synthesis, but instead require arginine uptake to cope with inhibition of *de novo* arginine synthesis. (A) RTqPCR analysis for *Sdc1* in mPDAC cells infected with lentiviruses encoding a dox inducible *Sdc1* targeting shRNA or an empty vector (EV) control with and without treatment with 1 µg/mL doxycycline or vehicle (n=2). (B) Macropinocytosis activity measured by kinetic DQ-BSA uptake and catabolism in mPDAC1-TIFM cells from (A). (C) Macropinocytosis activity measured by kinetic DQ-BSA uptake and catabolism in mPDAC1-TIFM cells treated with 10µM hydroxychloroquine (HQ) or vehicle (water). (D) Cell proliferation rate of mPDAC1-TIFM cells in same conditions as (C) (n=3). (E) Macropinocytosis activity measured by DQ-BSA uptake and catabolism in mPDAC1-TIFM cells cultured in TIFM with or without TIF concentrations of citrulline and ornithine. (F) Relative intracellular arginine levels of mPDAC1-TIFM cells grown in TIFM with or without PDAC IF concentrations of citrulline, ornithine, or arginine, as indicated (n=3). (G) Proliferation rate of mPDAC2-TIFM and mPDAC3-TIFM cells grown in TIFM with or without PDAC IF concentrations of citrulline, ornithine, or arginine, as indicated (n=3). For all panels, bar graphs represent the mean and the error bars represent ± SD.

**Figure 1 Source Data 1** Table of differentially expressed genes between mPDAC3-TIFM cells cultured in RPMI and isolated from orthotopic tumors generated by limma analysis of transcriptomic data as show in Fig. 1F.

**Figure 1 Source Data 2** Table of the top 40 differentially enriched MSigDB curated gene sets between mPDAC3-TIFM cells cultured in TIFM and RPMI as shown in Fig. 1H. Enrichment scores and significance values for the same gene sets are also shown for transcriptomic data of mPDAC3-TIFM cells isolated from orthotopic tumors and cultured in RPMI.

**Figure 1 Source Data 3** Table of GO gene set enrichment values comparing transcriptional differences between mPDAC3-TIFM cells cultured in TIFM and isolated from orthotopic tumors as show in Fig. 1I.

**Figure 2 Source Data 1** Table of metabolite consumption/release rates for mPDAC-TIFM cultured in TIFM and mPDAC1-RPMI cells cultured in RPMI.

**Figure 2 Source Data 2** Mass isotopomer distributions for all metabolites analyzed by LC-MS in Figure 2J.

**Figure 2 Source Data 3** All metabolite concentrations, tumor volumes, weights and densities measured in Fig. 2K used to calculate IF and intratumoral concentrations of arginine.

**Figure 4 Source Data 1** All metabolite concentrations measured in the IF of orthotopic mPDAC3-TIFM tumors from LysM-Cre^+/+^; Arg1^fl/fl^ and Arg1^fl/fl^ littermate controls in Fig. 4E. All metabolite concentrations measured in the IF of mPDAC3-TIFM orthotopic tumors after treatment with 100mg/kg of arginase inhibitor CB-1158 or vehicle in Fig. 4F.

**Supplementary File 1** Table with complete formulation of TIFM including all commercial suppliers and manufacturer part numbers for each metabolite included in TIFM.

**Supplementary File 2** Table of primers used for qPCR analyses and genotyping analysis of LysM-Cre and Arg1^fl/fl^ alleles. Table of shRNA hairpin and sgRNA sequences used in this study.

## Materials and Methods

### Formulation of Tumor Interstitial Fluid Media

TIFM is composed of 115 nutrients (Supplementary File 1) at levels that match the average of measurements in the IF of murine *LSL-Kras^G12D/+^; Trp53^flox/flox^ Pdx1-Cre* (KP^-/-^C) PDAC tumors (Sullivan et al., 2019a). The medium is composed of 10 pools of metabolites each of which is formulated by compounding dry powders of nutrients at appropriate ratios using a knife mill homogenizer. To generate the complete medium, the 10 metabolite mixture powders are added together and reconstituted in water with 10% dialyzed fetal bovine serum (FBS) to provide essential lipids, proteins and growth factors. The electrolytes provided in pool 3 are adjusted so that the electrolyte balance will be the same as RPMI-1640 medium, correcting for the sodium chloride in the FBS and counter ions in the various metabolites in TIFM. We performed quantitative LC-MS metabolite profiling (see **Quantification of metabolite levels in cell culture media**) to ensure concentrations of nutrients in TIFM are reproducibly close to the formulated concentration (Fig. 1B).

### Quantification of metabolite levels in cell culture media

For quantification of metabolites in cell culture media, quantitative metabolite profiling of fluid samples was performed on tissue culture media samples as previously described (Sullivan et al., 2019a). Briefly, chemical standard libraries of 149 metabolites in seven pooled libraries were prepared and serially diluted in HPLC grade water from in a dilution series from 5mM to 1µM to generate ‘external standard pools’, which are used for calibration of isotopically labeled internal standards and to quantitate concentrations of metabolites where internal standards were not available.

We then extracted metabolites from 5µL of either cell culture media samples or external standard pool dilutions using 45µL of a 75:25:0.1 HPLC grade acetonitrile:methanol:formic acid extraction mix with the following labelled stable isotope internal standards:

- ^13^C labeled yeast extract (Cambridge Isotope Laboratory, Andover, MA, ISO1)
- ^2^H_9_ choline (Cambridge Isotope Laboratory,Andover, MA, DLM-549)
- ^13^C_4_ 3-hydroxybutyrate (Cambridge Isotope Laboratory, Andover, MA, CLM-3853)
- ^13^C_6_ ^15^N_2_ cystine (Cambridge Isotope Laboratory, Andover, MA, CNLM4244)
- ^13^C_3_ lactate (Sigma Aldrich, Darmstadt, Germany, 485926)
- ^13^C_6_ glucose (Cambridge Isotope Laboratory, Andover, MA, CLM-1396)
- ^13^C_3_ serine (Cam-bridge Isotope Laboratory, Andover, MA, CLM-1574)
- ^13^C_2_ glycine (Cambridge Isotope Laboratory, Andover, MA, CLM-1017)
- ^13^C_5_ hypoxanthine (Cambridge Isotope Laboratory, Andover, MA, CLM8042)
- ^13^C_2_15N taurine (Cambridge Isotope Laboratory, Andover, MA, CNLM-10253)
- ^13^C_3_ glycerol (Cambridge Isotope Laboratory, Andover, MA, CLM-1510)
- ^2^H_3_ creatinine (Cambridge Isotope Laboratory, Andover, MA, DLM-3653)

Samples in extraction mix were vortexed for 10 min at 4°C and centrifugated at 15,000x rpm for 10 min at 4°C to pellet insoluble material. 20µL of the soluble polar metabolite supernatant was moved to sample vials for analysis by LC-MS as previously described (Sullivan et al., 2019b, 2019a).

Once LC-MS analysis was performed, XCalibur 2.2 software (Thermo Fisher Scientific) was used for metabolite identification. External standard libraries were used to confirm the m/z and retention time for each metabolite. For quantitative analysis, when internal standards were available, external standard libraries were used to quantitate concentrations of stable isotope labeled internal standards in the extraction mix. Once internal standard concentrations were obtained, peak area of an unlabeled metabolite in in the media samples was compared with the peak area of the quantified internal standard to determine the concentration of the unlabeled metabolite.

For metabolites for which an internal standard was not present in the extraction mix, external standard libraries were used to perform analysis of relevant metabolite concentrations. Briefly, the peak area of the metabolite was normalized to the peak area of a labelled stable isotope internal standard with the closest elution time, both in media samples and external standard library dilutions. Using the external standard library dilutions, we created a standard curve based on the linear relationship of the normalized peak area and the concentration of the metabolite, excluding those metabolites with an r^2^< 0.95. This standard curve was then used to interpolate the concentration of the metabolite in the media sample.

### Cell Isolation from tumors

Murine cancer cell lines were derived from tumor bearing C57Bl6J *LSL-Kras^G12D/+^; Trp53^flox/flox^; LSL-YFP; Pdx1-Cre* (KP^-/-^CY) mice to allow for fluorescent lineage tracing and isolation of cancer cells (Li et al., 2018). To isolate cancer cells from these tumors, the tumors were chopped finely and digested with 30mg/mL dipase II (Roche 28405100), 10mg/mL collagenase I (Worthington LS004194) and 10mg/mL DNase by constant rotation at 37C for 30 min. Digestion was quenched with 0.5M Ethylenediaminetetraacetic acid (EDTA) and cells were passed through a 70µM filter and rinsed with PBS before platting in RPMI-1640 (Corning 50–020-PC). YFP+ Cancer cells from each tumor were sorted twice on a BD FACSAria II Cell Sorter.

### Cell lines and cell culture

Use of cancer cell lines was approved by the Institutional Biosafety Committee (IBC no. 1560). All cell lines were tested quarterly for mycoplasma using the MycoAlert Mycoplasma Detection Kit (Lonza LT07-318). All cells were cultured in Heracell vios 160i incubators (Thermofisher) at 37°C and 5% CO2. Cell lines were routinely maintained in RPMI-1640 or TIFM supplemented with 10% diaFBS (Gibco, #26400-044, Lot#2244935P).

All cell culture was performed in static culture conditions. TIFM contains substantially lower levels of nutrients than most standard media formulations. Therefore, to ensure that there was not nutrient deprivation in static cultures the following modifications to standard tissue culture practices were made. Cells were cultured in larger volumes of media (8mL/35mm diameter well) to prevent depletion of nutrients during the culture. Additionally, media were replaced every 24 hrs. We routinely measured concentration of the most rapidly consumed nutrient, glucose, using a GlucCell glucometer (Bechard et al., 2020) to ensure that cultures used in experiments did not experience a greater than 30% drop in glucose availability, which is within the range of KP^-/-^C IF glucose concentration measurements (Sullivan et al., 2019a). Lastly, passaging TIFM maintained cells using standard trypsin (0.025%)/EDTA solution to detach cells leads to loss of viability upon replating of cells. Therefore, cells were detached with a 1:1 mixture of 0.5% trypsin-EDTA (Thermofisher) and serum free RPMI-1640 media (Thermofisher). This allowed routine passaging and plating of cells with less loss of viability. These modifications were followed for both TIFM and RPMI cultured cells.

### Determining cellular proliferation rate

Quantification of cellular proliferation rate was performed by sulforhodamine B (SRB) assay as described (Lien et al., 2021). Briefly, 10,000-15,000 cells were plated in 12-well plates in triplicates for each condition and allowed to attach overnight. After attachment, one set of triplicate wells were fixed by adding 10% trichloroacetic acid (TCA) to the media and incubating plates in 4°C to provide an ‘initial day’ value. Media was changed on remaining cultures and were allowed to grow for the indicated number of days. At the end of the growth period, cells were fixed by adding 10% trichloroacetic acid (TCA) to the media and incubating plates in 4°C for at least one hour. All wells, including the initial day wells were washed with deionized water, air-dried at room temperature, and dyed with SRB in 1% acetic acid for 30 min. After, cells were washed with 1% acetic acid three times and dried at 30°C for 15 minutes. SRB dye was solubilized with 10mM Tris pH 10.5 by gentle horizontal shaking for 5 min. Absorbance (abs) was measured at 510 nm in a clear 96-well plate using a BioTek Cytation 1 Cell Imaging Multi-Mode Reader. After all measurements were normalized to an averaged blank measurement (wells without cells but with media), growth rate was calculated using the following equation:

Doublings/day = log_2_(Final Day Abs_510_/Initial Day Abs_510_) / number of days elapsed in culture period

### Consumption/Release (Co/Re) analysis

Measurement of cellular consumption and release of metabolites was performed according to previous publications (Hosios et al., 2016; Jain et al., 2012; Muir et al., 2017). For the experimental set up, 100,000-150,000 cells were seeded in 2mL of culture medium in six-well plates with 3 technical replicates per condition per time point and allowed to attach overnight. The following day (day 1), cells were washed twice with 2mL PBS. They were then given 2mL of media, either TIFM or RPMI. An unspent media sample was collected at this time as well and stored at -80 °C. Cell number on day 1 was measured using a Vi-CELL XR Cell Viability Analyzer (Beckman Coulter). 24h later (day 2), 1mL of spent media from cells was collected, centrifuged and stored at -80 °C. Cell number was counted again. Quantification of metabolite levels in unspent (day 1) and day 2 cell culture media samples was performed as described in **Quantification of metabolite levels in cell culture media**.

To calculate Co/Re rates of a given metabolite, cell numbers on day 1 and day 2 were used to fit an exponential growth function, which integrated yielded the number of (cell · days) the day 2 media was conditioned. Changes in nutrient concentration in cultures were then normalized to this integrated growth curve to yield metabolite Co/Re per cell per unit of time.

### Experimental set up for consumption of arginine by GC-MS analysis

Cells were plated as described for consumption/release (Co/Re) analysis as described in **Consumption/Release (Co/Re) analysis**. The following day, cells were changed into either TIFM or TIFM without citrulline and ornithine. Both media were supplemented with 20µM extracellular arginine. Day 1 and day 2 media samples were collected and cell numbers were measured as in **Consumption/Release (Co/Re) analysis**.

10µL of each media sample were mixed 10µL of water containing ^13^C_6_,^15^N_4_ arginine at 20µM and 600µL cold HPLC grade methanol. The solution was then vortexed for 10 min, and centrifuged at 21,000xg for 10 min. Finally, 450μL of each extract was aliquoted, dried under nitrogen gas and stored at -80°C before further analysis. Sample derivatization GC-MS was then used to measure the arginine concentration in each media sample as described below in **GC-MS analysis of arginine**.

### RNA extraction, library preparation and transcriptomic analyses

#### Isolation of cultured and tumor cancer cell samples

mPDAC3-TIFM cells were plated at 200,000 (TIFM) to 350,000 (RPMI) cells per 6cm plate in triplicate cultures. RNA was extracted from cells 24 hrs later when the cells were proliferating exponentially, the cells were trypsinized and isolated by fluorescence activated cell sorting (FACS) for RNA extraction. For the *in vivo*, condition cells were isolated by FACS from end stage orthotopic mPDAC3-TIFM tumors, as described in **Cell Isolation from tumors**.

#### RNA extraction

Cells from all conditions were sorted by FACS prior to RNA extraction to eliminate the FACS sorting process as a confounder between cultured mPDAC3-TIFM cells and those isolated from orthotopic tumors. For FACS sorting, cells were stained with DAPI (750 ng/mL) to separate dead/dying cells from live cells, and live YFP+/DAPI-cells were sorted with a BD FACSAria II Cell Sorter with a 100µm nozzle directly into Qiagen RLT RNA extraction buffer. The ratio of RNA extraction buffer to sorted cellular volume was kept at 100µL of sorted sample per 350µL of RNA extraction buffer. Total messenger RNA (mRNA) was extracted using the RNeasy Micro Kit (Qiagen #74004) and RNA quality and quantity was assessed using the 2100 Bioanalyzer System (Agilent).

#### Library preparation and sequencing

Strand-specific RNA-SEQ libraries were prepared using an TruSEQ mRNA RNA-SEQ library protocol (Illumina). Library quality and quantity was assessed using the Agilent bio-analyzer and libraries were sequenced using an Illumina NovaSEQ6000.

#### Transcriptomic analyses

Data processing and analysis was done using the R-based Galaxy platform (https://usegalaxy.org/) (Afgan et al., 2018). Quality control was performed prior and after concatenation of the raw data with the tools *MultiQC* and *FastQC* respectively. All samples passed the quality check with most showing ∼20% sequence duplication, sequence alignment greater or equal to 80%, below and below 50% GC coverage, all of which is acceptable and/or indicative of good quality for RNASeq samples (Dündar et al., 2015; Parekh et al., 2016). Samples were then aligned, and counts were generated using the tools *HISAT2* (Galaxy Version 2.2.1+galaxy0, NCBI genome build GRCm38/mm10) and *featureCounts* (Galaxy Version 2.0.1+galaxy1), respectively. Differential expression analyses were performed with *limma* (Galaxy Version 3.48.0+galaxy1) and Genome Set Enrichment Analysis (GSEA) with *fgsea* (Galaxy Version 1.8.0+galaxy1) or GSEAPreranked (v6.0.12, https://gsea-msigdb.github.io/gseapreranked-gpmodule/v6/index.html) (Jain et al., 2012; Reich et al., 2006; Subramanian et al., 2005). GSEA plots were generated as previously described (Morris et al., 2019).

### Immunoblot analysis

For immunoblotting analysis, cells growing in log phase in a 6 well dish were washed with 2mL of PBS and lysed in 100μL RIPA buffer [25 mM Tris-Cl, 150 mM NaCl, 0.5% sodium deoxycholate, 1% Triton X-100, 1x cOmplete protease inhibitor (Roche)]. Cells were scraped and the resulting lysate was clarified by centrifugation at 21,000xg for 10 min. Protein concentration of the lysate was determined by BCA assay (Thermofisher). Proteins (20–30μg) were resolved on SDS-PAGE, 4 to 12% Bis-Tris Gels (Invitrogen) and transferred to a polyvinylidene difluoride membrane using the iBlot 2 Dry Blotting System (Invitrogen). Membrane was blocked with Intercept Blocking Buffer (Li-cor) at room temperature for 2h, stained with primary and secondary antibodies and then visualized using a LI-COR imager with Image Studio software version 2.1.10.

The following primary antibodies were used: Ass1 (Atlas HPA020896; 1:200 dilution), Vinculin (Proteintech 66305-1-lg; 1:10000 dilution), Beta-Actin (Proteintech 660009-1-lg; 1:10000 dilution). The following secondary antibodies: IRDye 680LT Goat Anti-Mouse Ig (Li-cor G926-68020; 1:10000 dilution) IRDye 800CW Goat anti-Rabbit IgG (Li-cor 926-32211; 1:10000 dilution), IRDye 800CW Goat anti-Mouse IgG (Li-cor 926-32210; 1:10000 dilution)

### qRT-PCR analysis

RNA was extracted using the RNeasy Mini Kit and optional on-the-column DNAse digestion (Qiagen). Extracted RNA was converted to cDNA by reverse transcription using the High-Capacity cDNA Reverse Transcription Kit (Applied Biosystems). Expression levels of *Sdc1* transcript were amplified using PowerUp SYBR Green Master Mix (Invitrogen) and custom primers (Supplementary File 2). Quantification was performed using a QuantStudio 3 Real-Time PCR System (Applied Biosystems). The average change in threshold cycle (ΔCt) values was determined for each of the samples relative to *Gapdh* levels and compared with vehicle control (ΔΔCt). Finally relative gene expression was calculated as (2^-ΔΔCt^). Experiments were performed in with triplicate cultures and RNA extraction.

### GC-MS analysis of arginine

Dry polar metabolites extracts from intracellular extracts or media samples were derivatized with 16μL MOX reagent (ThermoFisher) for 1h at 37°C and then with 20μL 1% tert-butyldimethylchlorosilane in N-tert-Butyldimethylsilyl-N-methyltrifluoroacetamide (Sigma Aldrich) for 3h at 60°C. Derivatized samples were analyzed with an 8890 gas chromatograph system (Agilent Technologies) with a HP-5ms Ultra Inert column (Agilent Technologies) coupled with an 5997B Mass Selective Detector (MSD) mass spectrometer (Agilent Technologies). Helium was used as the carrier gas at a flow rate of 1.2 mL/min. One microliter of sample was injected in splitless mode at 280°C. After injection, the GC oven was held at 100°C for 1 min. and increased to 300°C at 3.5 °C/min. The oven was then ramped to 320°C at 20 °C/min. and held for 5 min. at this 320°C. The MS system operated under electron impact ionization at 70 eV and the MS source was held at 230 °C and quadrupole at 150 °C. Detector was set in scanning mode with a scanned ion range of 100–650 m/z. Mass isotopomer distribution was determined using ion fragments integration for each individual metabolite (Lewis et al., 2014) and natural abundance correction was performed using IsoCor (Millard et al., 2019).

### Isotopic labeling experiments in cell culture and intracellular metabolite extraction

To measure steady state labeling of polar metabolites by citrulline in cultured cells, triplicate cultures of 150,000 cells/well were seeded in a 6 well dish in 2 mL of medium. Cells were allowed to attach overnight. The following day cells were washed twice with PBS and then incubated with 8mL for 8 or 24h in TIFM with ^13^C_5_-citrulline (Cambridge Isotope Laboratories, CLM-8653) added at TIFM concentrations. Immediately after the labeling period, cells were quickly washed with ∼8mL of ice-cold blood bank saline. Cellular metabolites were extracted with addition of 600μL of an ice-cold methanol followed by scraping the cells on ice. The solution was then vortexed for 10 min, and centrifuged at 21,000xg for 10 min. 450μL of each extract was aliquoted to fresh sample tubes, dried under nitrogen gas and stored at -80°C before further analysis.

### CRISPR knockout and re-expression of *Ass1*

sgRNAs targeting *Ass1* (Supplementary File 2) were generated through the Broad Institute’s Genetic Perturbation Platform Web Portal (https://portals.broadinstitute.org/gpp/public/). Oligonucleotide pairs were manufactured by Integrated DNA Technologies (IDT) and cloned into lentiCRISPRv2 (Addgene: #52961) as previously described (Sanjana et al., 2014; Shalem et al., 2014). HEK293T cells (Dharmacon) were transfected with the *Ass1* targeting lentiCRISPRv2 vectors and the lentiviral packing plasmids psPAX2 (Addgene: #12260) and pMD2.G (Addgene: #12259). Medium was replaced after 24h, and lentivirus was harvested after 48h. Subconfluent mPDAC3-TIFM cells were infected with lentivirus using 8µg/mL polybrene and infected cells were selected in 2µg/mL puromycin and maintained with 100µM arginine. Single cell clones with immunoblot-confirmed loss of Ass1 were selected. A single cell clone without detectable Ass1 expression was transformed with a lentivirus produced as above with a vector encoding CMV-driven murine *Ass1* cDNA that would not be targeted by the *Ass1* sgRNA (VectorBuilder).

### shRNA knockdown of *Sdc1*

Hairpin sequences targeting *Sdc1* were obtained from (Fellmann et al., 2013). Oligonucleotide pairs were manufactured by IDT and cloned into a lentiviral LT3GEPIR vector (Addgene: #111177) to allow for doxycycline-inducible repression of gene expression. Lentiviral transfection and transformation were performed as described in **CRISPR knockout and re-expression of *Ass1*** and successfully transformed cells were selected and maintained with 2μg/mL puromycin. Cells transformed with LT3GEPIR without an shRNA were used as a control.

### Analysis macropinocytic capacity by DQ-BSA

The macropinocytic capacity of PDAC cells was assessed using a DQ Red BSA (Invitrogen) uptake assay. Cells were seeded at either 15,000 cells/well for 12 wells or 50,000 cells/well for 6 wells and allowed to attach over night. The following day the media was replaced with fresh media + 0.02mg/mL of the DQ Red BSA fluorogenic substrate and cells were harvested at different timepoints for up to 6 hours. Cells were then washed with PBS, trypsinized, washed again with PBS, fixed in 4% paraformaldehyde for 15 minutes at 4°C and DQ Red BSA fluorescence was quantified by flow cytometry in at least 10,000 cells per sample.

### Animal Experiments

Animal experiments were approved by the Institutional Animal Care and Use Committee (IACUC, Protocol #72587) and performed in strict accordance with the Guide for the Care and Use of Laboratory Animals of the National Institutes of Health (Bethesda, MD). Mice were housed in a pathogen-free animal facility at the University of Chicago with a 12 h light/12 h dark cycle, 30–70% humidity and 68–74°F temperatures maintained.

#### Orthotopic tumor implantation and monitoring

C57BL6J mice 8-12 weeks of age were purchased from Jackson Laboratories. 250,000 cells/tumor were resuspended in 20µL of 5.6mg/mL Cultrex Reduced Growth Factor Basement Membrane Extract (RGF BME; R&D Biosystems #3433-010-01) and serum-free RPMI (SF-RPMI) solution. The BME:cellular mixture was injected into the splenic lobe of the pancreas of the mice as previously described (Erstad et al., 2018). After implantation mice, were monitored daily by abdominal palpation.

#### In vivo Arg1 knockout

LysM-Cre and Arg1^fl/fl^ C57BL6J mice were bred to generate LysM-Cre ^+/+^; Arg1^fl/fl^ and litter mate control Arg1^fl/fl^ mice. All mice were genotyped using primes described in Supplementary File 2. Animal husbandry was carried out in strict accordance with the University of Chicago Animal Resource Center guidelines. Tumor implantation as described above was performed in mice at 8-12 weeks of age.

#### In vivo arginase-1 pharmacological inhibition

Orthotopic tumors were implanted in C57BL6J mice at 8-12 weeks of age. 4 weeks after induction, once tumors were close end stage disease, CB-1158 (MedChem Express) dissolved in sterile water was administered by oral gavage at 100mg/kg as previously described (Steggerda et al., 2017). The acidity caused by the HCl in the drug solution was neutralized by adding equivalent amount of NaOH and an equivalent NaCl in sterile water solution was prepared as vehicle. 2hrs after treatment with CB-1158 or vehicle, mice were euthanized by cervical dislocation, and tumors were harvested for TIF extraction.

#### In vivo ^15^N_2_-glutamine tracing

Orthotopic tumors were implanted in C57BL6J mice at 8-12 weeks of age. 4 weeks after induction tumor-bearing mice and healthy littermate controls were treated with ^15^N_2_-glutamine (Cambridge Isotope Laboratory #NLM-1328-PK) dissolved in sterile phosphate buffered saline at 7.2mg/animal by tail vein injection as previously described (Lane et al., 2015). Briefly, animals were dosed three times at 15-minute intervals. 15 minutes after the final dose, ∼100uL of blood were be obtained by submandibular sampling as described previously (Parasuraman et al., 2010) and animals were euthanized. The tumor or pancreas from each animal was then harvested and immediately snap frozen using a BioSqueezer (BioSpec) cooled with liquid nitrogen and stored at -80°F until further analysis.

#### IF isolation from PDAC tumors

IF was isolated from tumors as described before (Sullivan et al., 2019a). Briefly, tumors were rapidly dissected after euthanizing animals. Tumors were weighed and rinsed in blood bank saline solution (150 mM NaCl) and blotted on filter paper (VWR, Radnor, PA, 28298–020). The process of dissection and tumor preparation took < 3min. Tumors were cut in half and put onto 20µm nylon mesh filters (Spectrum Labs, Waltham, MA, 148134) on top of 50 mL conical tubes, and centrifuged for 10min. at 4°C at 400xg. IF was then collected, snap-frozen in liquid nitrogen and stored at -80°C until further analysis.

### HPLC-MS-MS analysis amino acid levels in PDAC IF samples upon arginase inhibition

IF samples were analyzed by High-Performance Liquid Chromatography and Tandem Mass Spectrometry (HPLC-MS-MS). Specifically, the system consisted of a Thermo Q-Exactive in line with an electrospray source and an Ultimate3000 (Thermo) series HPLC consisting of a binary pump, degasser, and auto-sampler outfitted with a Xbridge Amide column (Waters; dimensions of 3.0 mm × 100 mm and a 3.5 µm particle size). The mobile phase A contained 95% (vol/vol) water, 5% (vol/vol) acetonitrile, 10 mM ammonium hydroxide, 10 mM ammonium acetate, pH = 9.0; B was 100% Acetonitrile. The gradient was as following: 0 min, 15% A; 2.5 min, 64% A; 12.4 min, 40% A; 12.5 min, 30% A; 12.5-14 min, 30% A; 14-21 min, 15% A with a flow rate of 150 μL/min. The capillary of the ESI source was set to 275 °C, with sheath gas at 35 arbitrary units, auxiliary gas at 5 arbitrary units and the spray voltage at 4.0 kV. In positive/negative polarity switching mode, an m/z scan range from 60 to 900 was chosen and MS1 data was collected at a resolution of 70,000. The automatic gain control (AGC) target was set at 1 × 106 and the maximum injection time was 200 ms. The targeted ions were subsequently fragmented, using the higher energy collisional dissociation (HCD) cell set to 30% normalized collision energy in MS2 at a resolution power of 17,500. Besides matching m/z, target metabolites are identified by matching either retention time with analytical standards and/or MS2 fragmentation pattern. Data acquisition and analysis were carried out by Xcalibur 4.1 software and Tracefinder 4.1 software, respectively (both from Thermo Fisher Scientific).

### Measurement of intratumoral metabolite levels and metabolite isotope labeling patterns

Cryogenically frozen tumor pieces were ground to a fine homogenous powder with a liquid nitrogen cooled mortar and pestle. ∼30mg of tissue powder was weighed into sample tubes, and metabolites were extracted with 600µL HPLC grade methanol, 300µL HPLC grade water, and 400µL chloroform. Samples were vortexed for 10min at 4°C, centrifuged 21,000xg at 4°C for 10 min. 400µL of the aqueous top layer was removed into a new tube and dried under nitrogen. Dried tumor extracts were resuspended in 100µL HPLC grade water and LC-MS analysis was performed as described before(Sullivan et al., 2019b, 2019a). XCalibur 2.2 software (Thermo Fisher Scientific) was used identification and relative quantification for metabolites. Natural abundance correction was performed using the IsoCor (Millard et al., 2019).

### Measuring intratumor and IF concentrations of amino acids

To quantitatively measure IF amino acid abundance, polar metabolites were extracted from 5µL IF samples using 45µL 75:25:0.1 HPLC grade acetonitrile:methanol:formic acid extraction mix into which a mixture of isotopically labeled amino acids of known concentrations (Cambridge Isotope Laboratories, MSK-A2-1.2) was added. Samples were vortexed for 10 min, and centrifuged at maximum speed for 10 min. 30 μL of each extract was removed and dried under nitrogen gas and stored −80°C until further analysis by LC-MS as described in **Quantification of metabolite levels in cell culture media**. Amino acids amounts in IF samples were determined by comparison of peak areas of unlabeled amino acids with peak areas of labeled amino acids that were present at known amounts. Concentrations were determined by dividing the amino acid amount by the 5µL volume of the IF sample.

To measure amino acid amounts in intratumoral samples, intratumoral metabolites were extracted from ∼30mg and dried down in **Measurement of intratumoral metabolite levels and metabolite isotope labeling patterns**. Dried samples were rehydrated with 2:1 methanol:water into which a mixture of isotopically labeled amino acids of known concentrations (Cambridge Isotope Laboratories, MSK-A2-1.2) was added. Samples were then analyzed by LC-MS as described in **Quantification of metabolite levels in cell culture media**. Amino acid amounts per tumor weight were determined by comparison of peak areas of unlabeled amino acids with peak areas of labeled amino acids that were present at known amounts and dividing by the mass of tumor extracted.

To compare metabolite concentrations between tumor and TIF samples, the density for orthotopic mPDAC tumors was needed to convert amino acid amount per unit tumor mass into a concentration (amino acid amount per unit volume). The density of freshly isolated mPDAC3-RPMI tumors was measured by measuring tumor mass and calculating the volume (V) of the tumors with the following formula:

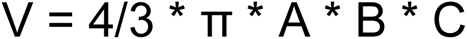

where A, B, C are the lengths of the semi-axes of an ellipsoidal shape, which were measured from tumors with an electronic caliper (Thermofisher). Tumor density was then calculated by dividing the tumor mass by the calculated volume. Tumor density was then used to convert amino acid amount per tumor mass measurements into an intratumoral concentration.

### Human samples regulation

Human histology samples were obtained under approval by the Institutional Review Boards at the University of Chicago (IRB 17-0437).

### Immunohistochemistry

For ARG1 and ASS1 staining, the slides were stained using Leica Bond RX automatic stainer. Dewax (AR9222, Leica Microsystems) and rehydration procedure were performed in the system and a 20 min treatment of epitope retrieval solution I (Leica Biosystems, AR9961) was applied. anti-Arginase-1 (1:100, Cell Signaling #93668) or anti-Ass1 (1:100, Atlas HPA020896;) and were applied on tissue sections for 60min. Antigen-antibody binding was detected using Bond polymer refine detection (Leica Biosystems, DS9800). The tissue sections were counter stained with hematoxylin and covered with cover glasses.

For F4/80 staining, tissue sections were deparaffinized and rehydrated with xylenes and serial dilutions of EtOH to deionized water. They were incubated in antigen retrieval buffer (DAKO, S1699) and heated in steamer at 97°C for 20 minutes. Anti-mouse F4/80 antibody (1:200, MCA497GA, AbD Serotec) was applied on tissue sections for 1hr at room temperature. Tissue sections were washed with Tris buffered saline and then incubated with biotinylated anti-rat IgG (10 μg/ml, BA-4001, Vector laboratories) for 30 min at room temperature. Antigen-antibody binding was detected by Elite kit (PK-6100, Vector Laboratories) and DAB (DAKO, K3468) system.

## Acknowledgements

We thank all members of the Muir lab for many discussions, feedback, and support. We thank Lev Becker and Catherine Reardon from the University of Chicago for the generous gift of LysM-Cre and Arg1fl/fl mice and guidance with the use of these animals. We thank Brandon Faubert for the useful discussions about in vivo isotope labelling experiments and critical review of the manuscript. We also thank Mark Sullivan and Laura Danai for discussions and feedback about the manuscript. We thank the Metabolite Profiling Core Facility at the Whitehead Institute for processing metabolomics samples and assistance with data analysis. We thank the Metabolomics Core Facility at Robert H. Lurie Comprehensive Cancer Center of Northwestern University for assistance with metabolomics services. We thank The University of Chicago Animal Resources Center (RRID:SCR_021806), especially Ani Solanki, for their assistance with animal models of pancreatic cancer. We thank The University of Chicago Genomics Facility (RRID:SCR_019196), Human Tissue Resource Center (RRID:SCR_019199) and Cytometry and Antibody Technology Facility (RRID: SCR_017760) for their invaluable technical assistance. All of these facilities receive financial support from the Cancer Center Support Grant (P30CA014599). We also thank resources at the University of Chicago dedicated to promoting the recruitment and retention and support of underrepresented minority (URM) students in science, including The Graduate Recruitment Initiative Team (GRIT), and those promoting accessible mental health resources, including UChicago Student Wellness. This work was supported by grants to AM from the American Cancer Society (IRG-16-222-56), the University of Chicago Cancer Center Support Grant (P30 CA14599), the Pancreatic Cancer Action Network (2020 Career Development Award), the Brinson Foundation, the Cancer Research Foundation and the Ludwig Center for Metastasis Research. KAF was supported by the National Cancer Institute (R01 CA200310). JJAS, PBJ and CS were supported by the NCI (T32 CA009594). JJAS also received support from the MaryEllen Connelan Award, the Robert C. and Mary Jane Gallo Scholarship Fund and the Harper Fellowship at the University of Chicago.

## Data availability

Sequencing data from Figures 1 and 3 have been deposited in GEO under accession code GSE199163: https://www.ncbi.nlm.nih.gov/geo/query/acc.cgi?acc=GSE199163. Source data files with measured metabolite concentrations and isotopic labeling patterns are provided for Figures 2 and 4 and we are in the process of depositing raw mass spectra from these experiments in Metabolomics Workbench. Prior to upload, this data is available upon request.

